# Adaptation of visual responses in degenerating *rd10* and healthy mouse retinas during ongoing electrical stimulation

**DOI:** 10.1101/2025.02.19.639073

**Authors:** Archana Jalligampala, Hamed Shabani, Eberhart Zrenner, Daniel L. Rathbun, Zohreh Hosseinzadeh

**Author notes:** Corresponding Authors: Archana Jalligampala, Daniel L. Rathbun.

## Abstract

**Objective:** Visual adaptation is a physiological and perceptual process by which the visual system adjusts to changes in the environment or visual stimuli. This process is fundamental to how we perceive the world around us and allows our visual system to efficiently encode and process visual information. The retina incorporates adaptaion with its dozens of functionally distinct retinal ganglion cell types. Meanwhile, the field of retinal prostheses is increasing its understanding of electrical adaptation and cell-specific stimulation. However, very little is known about the interaction of visual and electrical stimulation on the adaptation of retinal ganglion cell types.

**Methods/Approach:** Recording with a microelectrode array (MEA), we presented an ON and OFF full-field, visual stimulus to characterize various visual response parameters in healthy and degenerating *rd10* mouse retinas. We then evaluated visual response changes before and after blocks of monophasic voltage-controlled electrical pulse stimulation.

**Main Results:** A history of electrical stimulation strengthened visual responses in WT retina, even when changes attributable to *in vitro* visual adaptation were taken into account. In *rd10* retinas, electrical adaptation counteracted the baseline *in vitro* visual adaptation. In all cases, adaptation often affected the ON and OFF visual response components differentially. Consequently, the ON/OFF classification of indiviual cells changed as a result of adaptation.

**Significance:** Electrical stimulation-induced changes in the retina should be considered in the encoding of visual stimuli by retinal prosthetic devices. *In vitro* investigations for bionic vision should strive to probe electrical responsiveness after adaptation to ongoing electrical stimulation has achieved a steady-state.

## Introduction

The human visual system is a remarkable entity, capable of perceiving and adapting to a range of visual stimuli, from the dim glow of starlight to the brilliance of a sunny day (Webster 2015). Over the past century, many luminaries like Ramón y Cajal, Hartline, Barlow, Kuffler, Lettvin, Hubel, and Wiesel have collectively contributed to today’s understanding of information processing via the visual system (Kuffler 1953, Rodieck & Stone 1965, Hubel & Wiesel 1998, Hartline 1938, Barlow 1953, Lettvin *et al*. 1959). Central to the remarkable dynamism of the visual system is the retina’s capacity for adaptation – to recalibrate and fine-tune its responses to suit the ever-changing demands of the visual input. There is an ever growing body of literature illuminating the various visual adaptations occurring in the retina (Baccus & Meister 2002, Demb 2002, Moore *et al*. 2011, Tikidji-Hamburyan *et al*. 2015, Tikidji-Hamburyan *et al*. 2017, Borghuis *et al*. 2018).

The development of retinal prosthetic devices offers a promising solution to the vision loss experienced by individuals with retinal degenerative diseases such as retinitis pigmentosa and age-related macular degeneration (Dagnelie 2012, Ayton *et al*. 2020). However, the aforementioned concept of retinal adaptation takes on profound significance in the context of retinal prosthetics where adaptation may be a barrier to ongoing perception (Fernandez 2018). Many studies have also investigated electrical adaptation in the retina (Fried *et al*. 2006, Jensen & Rizzo 2007, Freeman & Ray 2010, Fried 2011, Lorach *et al*. 2015, Im & Fried 2016,). Our own lab has previously demonstrated that voltage tuning curves displayed substantial hysteresis due to adaptation to electrical stimulation occurring over many seconds (Hosseinzadeh *et al*. 2018). Despite these efforts, little is known about the complex interaction between electrical and visual adaptation.

An important limitation on adaptive mechanisms is the receptive field of a neuron. First introduced by Weber in the context of tactile stimulation as ‘sensory circles’ (Weber 1846), and renamed by Sherrington (1906). Receptive fields (RF), in essence, are the spatial domains within which a neuron or a group of neurons can detect and respond to sensory stimuli. These fields vary in size, shape, and functional properties across the visual system, from the retinal ganglion cells to the complex networks of the visual cortex (Hubel & Wiesel, 1962). The history of the visual RF can be traced back to 1938, from the works of Hartline, who found that by stimulating a small circular area in the retina one could elicit an excitatory response in the optic nerve fiber of the frog (Hartline 1938). He termed this area the visual RF. Based on the responses, he classified the fibers as ON, OFF, and ON-OFF, which corresponded to responses to either light onset, light offset, or both, respectively. In 1953, Barlow and Kuffler established the existence of lateral inhibition in the frog and cat eye, respectively, by discovering the antagonistic center-surround organization of the RF (Barlow 1953, Kuffler 1953). Kuffler further classified the RF as *ON-center* RF (center activated by light onset) and *OFF-center* RF (center activated by light offset). Furthermore, they extended this work to classify the response durations of these RFs as *sustained* and *transient* under specific conditions of stimulus size and state of adaptation. Shortly after, Lettvin and Hubel & Wiesel in their iconic works classified visual cortex RFs selective to different complex features (shape, size, orientation, the position of the stimulus relative to the background, movement, and ocularity) (Letvinn 1959, Hubel & Wiesel 1959).

A recent study in mouse retina by Baden and co-workers (Baden *et al*. 2016) has shown that, beyond the simple ON, OFF, sustained, and transient response types, there are at least 30 different physiological types of retinal outputs. Other recent investigations into bionic vision as a cure for blindness have additionally shown that electrical input filters can vary according to retinal ganglion cell (RGC) type, suggesting the possibility for cell-specific stimulation (Sekhar *et al*. 2017, Ho *et al*. 2018). In more recent effort, we sought to compare the differences in prefered electrical input with this well-established catalog of dozens of RGC types, relying on their functional responses to light stimulation as a reference point (Shabani *et al*. 2021, Shabani *et al*. 2024). The present study provides insights into the adaptation of ON and OFF responses that occur when retinas receive electrical stimulation. Understanding such adaptation is an essential aspect of maximizing the effectiveness of bionic vision devices and improving the perceptual experience of recipient patients.

In this study, using ON and OFF full-field flash stimulus, we characterize the cells’ visual responses and evaluate how this response changes following electrical stimulation with voltage-controlled pulses. The results enhance our understanding of adaptive mechanisms and electrical stimlation of the retina.

## Materials and Methods

### Animals

The animals were housed under standard white cyclic lighting, mimicking regular daily rhythms. They had free and ample access to food and water. Adult wild-type C57Bl/6J (Jackson Laboratory, Bar Harbor, ME, USA) and *rd10* (on a C57Bl/6J background; Jackson Laboratory) strains were used, with age ranging from postnatal day 28 to 37 for both strains. The *rd10* strain is a well-established model for retinal degeneration in which the retina is unhealthy, but not yet blind at the ages examined. For each strain, three male and two female mice were used. For the external control condition, age-matched mice were used for both strains. For each strain of the control mice, two male mice were used. All procedures were approved by the Tübingen University committee on animal protection (Einrichtung für Tierschutz, Tierärztlichen Dienst und Labortierkunde directed by Dr. Franz Iglauer) and performed by the Association for Research in Vision and Ophthalmology (ARVO) statement for the use of animals in ophthalmic and visual research.

### Retinal Preparation

For dissecting the retina, the mice were anesthetized by CO_2_ inhalation. Following CO_2_ inhalation, the mice were checked for absence of withdrawal reflex by pinching the between-toe tissue and then euthanized by cervical dislocation. Under normal room lighting, the eyes were removed to carbogenated (95% O_2_ and 5% CO_2_) artificial cerebrospinal fluid (ACSF) solution containing the following (in mM): 125 NaCl, 3.5 KCl, 2 CaCl_2_, 1 MgCl_2_, 1.25 NaH_2_PO_4_, 25 NaHCO_3_, and 25 Glucose, pH 7.4. For each eye, the cornea, ora serrata, lens, and vitreous body were removed, the retina was detached from the pigment epithelium, and the optic nerve was cut at the base of the retina. Special care was taken to remove all traces of vitreous material from the inner surface of the retina to optimize contact between the nerve fiber layer and recording electrodes. The retinas were maintained in carbogenated ACSF until needed. For recording, a retinal half was mounted with the ganglion cell layer down on a planar microelectrode array (MEA). Two small paint brushes were used to orient and flatten the retinal half without risking damage to the MEA. A dialysis membrane (Cellu Sep, Membrane Filtration Products Inc., Seguin, Texas, USA) mounted on a custom Teflon ring was lowered onto the retina to press it into closer contact with the MEA (Meister *et al*. 1994). After securing the MEA under the preamplifier, the retina was continuously superfused with carbogenated ACSF (∼6 ml/min) maintained at 33° C using both a heating plate and a heated perfusion cannula (HE-Inv-8 & PH01; Multi Channel Systems, Reutlingen, Germany). A stabilization time of > 30 min was provided prior to recording the data.

### Microelectrode Array (MEA) and Data Acquisition

For recording the spiking responses from the RGCs, a planar MEA containing 59 circular titanium nitride electrodes (diameter: 30 µm, interelectrode spacing: 200 µm; Multi Channel Systems, Reutlingen, Germany) arrayed in an 8 x 8 rectilinear grid layout, and with Indium tin oxide (ITO) electrode tracks insulated by Silicon Nitride (Si_3_N_4_) on a glass substrate was used. Four electrodes were absent from the four corners of the grid, and one electrode was substituted with a large reference electrode. The impedances of the electrodes were approximately 200-250 kΩ at 1 kHz, measured using a nanoZ impedance meter (Plexon Inc., TX, USA) in saline. The MEA60 system (MCS, Reutlingen, Germany) was used for data acquisition including: the RS-232 interface, a 60-channel preamplifier with integrated filters and a blanking circuit (MEA 1060-Inv-BC) controlled by MEA_Select software to reduce recording noise by grounding any defective electrodes, and to assign electrical stimulation waveforms to the selected electrode. Data were collected using the MC_Rack program on a personal computer running Windows XP and fitted with MC_Card data acquisition hardware and an analog input card to record stimulus trigger signals. The raw data were recorded at a rate of 50 kHz/channel with a filter bandwidth ranging from 1 Hz – 3 kHz and amplification gain of 1100.

### Electrical Stimulation

In our previous study (refer to the Stimulation subsection of Material and Methods; Jalligampala *et al*. 2017), a detailed description of the electrical stimulation procedure is provided. Briefly, the stimulus pulses were generated using a stimulus generator (STG 2008, Multi Channel Systems, Reutlingen, Germany) and delivered from the ganglion cell side of the retina (epiretinally) via one of the 59 electrodes – which was always an interior electrode and chosen based on proximity to electrodes with robust neural signals to ensure a maximum number of recorded neurons. As the retinal network can be activated from either side of the retina by reversing the polarity of stimulation (Im & Fried 2015, Boinagrov *et al*. 2014, Eickenscheidt *et al*. 2012), and due to the ease of accessing the retina, the electrodes of the MEA were used to simultaneously stimulate and record from the ganglion cell (epiretinal) side of the flat-mounted retina. The stimulus (**Fig. 1a**, left panel) consisted of monophasic rectangular voltage pulses, each with one of the following amplitudes (0.3, 0.5, 1.0, 1.5, 2.0, 2.5 V), included both cathodic (-V) and anodic (+V) stimuli, and the following durations (60, 100, 200, 300, 500, 1000, 2000, 3000, 5000 µs). While 0.1 V was also presented, the stimulator was later found to be unable to deliver the waveform faithfully. Accordingly, this stimulus was excluded from analysis in the previous study. To reduce the possibility of electrolysis and electrode degradation only voltage/duration combinations that fell within safety, limits were delivered (Multi Channel Systems 2010). Within each increasing voltage block, the durations were presented in 5 sequential, uniquely randomized sets, for a total of 5 repetitions for each voltage/duration combination, with an interval of 5 s after each pulse to allow the recovery of RGC responsiveness. Additionally, for each experiment, the beginning polarity was randomly chosen and alternated with subsequent blocks after that. Before and after each stimulation block, spontaneous activity was recorded for ∼30 s.

**Figure 1:**
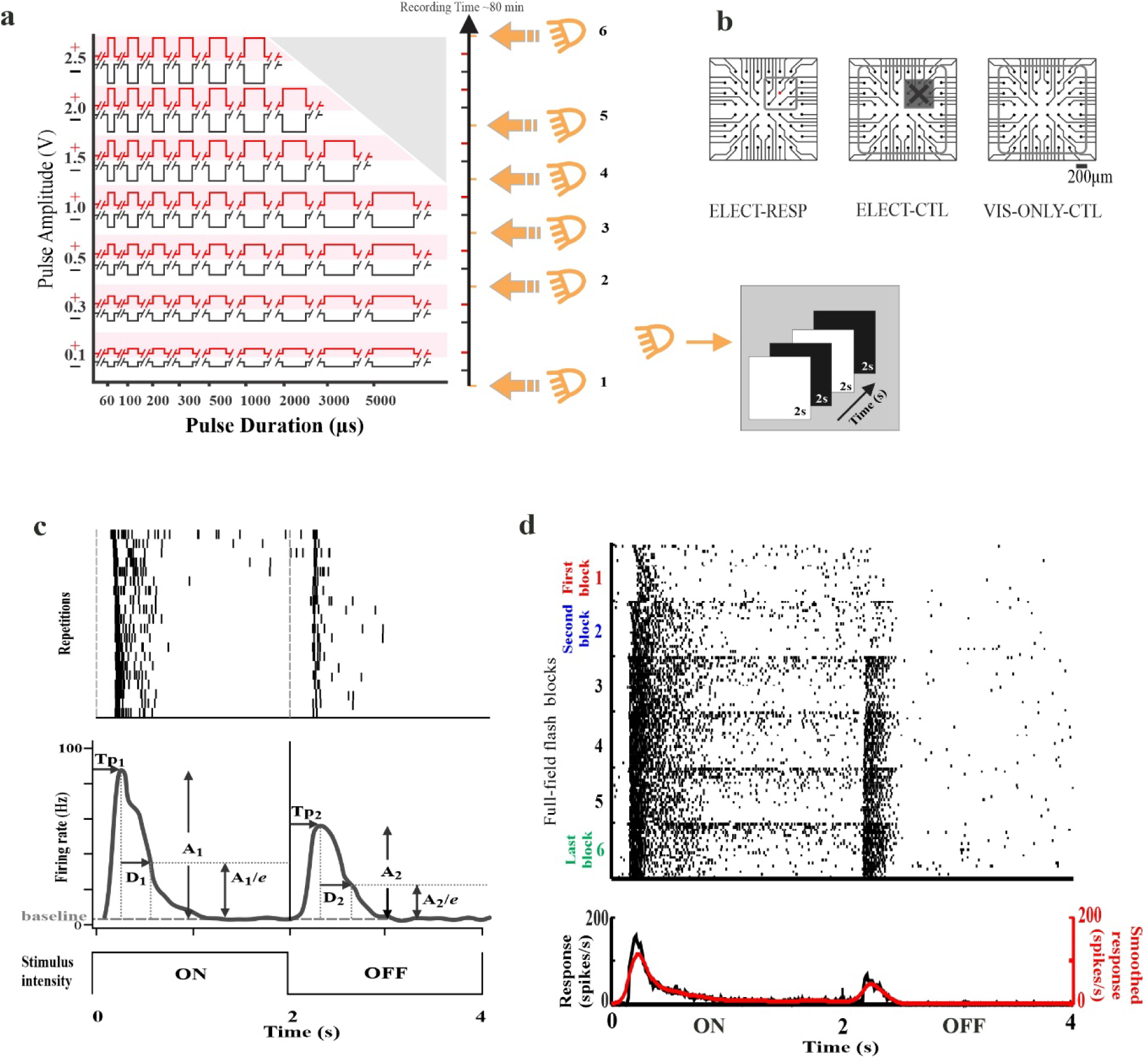
Experimental design. **(a)** Electrical stimuli were presented over a range of voltage/duration combinations. Constant-voltage stimulus blocks consisted of 5 repetitions of 9 durations. Within each repetition, the durations were randomized, 5 s separated pulses. Voltage blocks were presented from lowest to highest amplitude. Subsequent voltage blocks were separated by 150 s or greater. A block of full-field flash stimulus (20 repetitions of 2 s ON & 2 s OFF) was interleaved between voltage blocks throughout the experiment. Voltage-duration combinations that exceeded safety limits were omitted (grey triangle). **(b)** The different test (ELECT-RESP) and control conditions (internal ELECT-CTL and external VIS-ONLY-CTL). Red denotes stimulating electrode. Open grey box indicates electrodes on which included cells were recorded. Grey box with cross indicates that cells recorded on these 9 electrodes were excluded for the ELECT-CTL condition. **(c)** Flash response characterization according to Carcieri *et al*. 2003. *(top)* Spike rasters of a full-field flash stimulus (20 trials, 2 s ON 2 s OFF). *(middle)* Peristimulus time histogram (PSTH) derived from the rasters. *(bottom)* Visual stimulus time course. A_1_ and A_2_ are relative (to baseline) response amplitude for ON and OFF. Tp_1_ and Tp_2_ are time to peak (latency) for ON and OFF. D_1_ and D_2_ are durations for ON and OFF. **(d)** Visual response changes were evaluated by comparing First (red) to Second (blue) and First to Last (green) using the average PSTH binned at 2 ms intervals for all responses. Gaussian smoothing filter σ = 4 ms.

### Visual Stimulation

Visual stimuli were presented to the retina from below through the transparent MEA by a commercially available DLP-based projector (K10; Acer Inc., San Jose, California, USA). The image was focused and centered onto the plane of the retina directly over the MEA with a custom-built series of optical components and manual microdrive with three degrees of freedom. Visual stimuli were controlled with custom software. Each visual stimulus block consisted of a full-field (∼3 x 4 mm) ‘flash’ stimulus, cycling 2 s ON (40 klx) followed by 2 s OFF (20 lx), 20 times without pause (mean illuminance = 20 klx, 99.9% Michelson contrast; **Fig. 1a**, right panel). The brightness range chosen covered a wide range of intensities occurring in the natural environment (Rodieck 1998). The six visual stimulus blocks were interleaved before, after, and within an electrical stimulation experiment that spanned ∼80 min of recording time, including the First and Last flash blocks.

### Test and Control Conditions

Test Condition: The test condition was the visual response parameters obtained from RGCs which were responsive to electrical stimulation (see Jalligampala *et al*. 2017: Data analysis, responsive RGC inclusion criteria). Briefly, for a cell to be classified as electrically responsive, 1) at least three of the 96 responses (corresponding to the 96 unique electrical stimuli (Fig. 1a) was greater than two standard deviations (SD) above the average spontaneous rate (threshold), and 2) such responses had a firing rate equivalent to at least 8.89 Hz (at least four spikes within the 5 integration windows of 90 ms each, corresponding to the 5 presentations of that stimulus). An additional inclusion criterion was applied to the data set. Because electrical field strength falls with increasing inter-electrode distance from the stimulating electrode ̶ grouping responses from a wide range of distances adds unnecessary variability. Therefore, only responses from cells recorded at the 8 electrodes surrounding the stimulating electrode (distances of 200 µm and 283 µm) were included in the test condition. In referring to these cells that were near to the stimulating electrode (in red, Fig. 1b, left panel) and were electrically responsive we will use the term electrically responsive (ELECT-RESP).

#### Control Conditions

We compared the test condition to two different control conditions.

1) Internal control condition- As an internal control condition, we considered cells which were electrically nonresponsive, i.e., cells which were not well-driven by the electrical stimulation in the same recording (tissue). To minimize the effects of an electrically responsive network, we only considered nonresponsive cells at an inter-electrode distance > 300 µm from the stimulating electrode. A key feature of the internal control condition was that the responses from these cells were from the same tissue. Hence the factor of tissue variability is minimal. However, one caveat to this control condition is that electrical responses have been shown to extend at least as far as 800 µm from the stimulating electrode (Eickenscheidt *et al*. 2012, Ryu *et al*. 2009, Stett *et al*. 2007, Jalligampala *et al*. 2017, Wilke *et al*. 2011). In referring to these electrically nonresponsive cells recorded distant (> 300 µm) from the stimulating electrode we will use the term electrical control (ELECT-CTL, Fig. 1b, middle panel).
2) External control condition- A more appropriate control condition to strengthen our hypothesis was to stimulate the retinal tissue with only visual stimuli in the absence of any electrical stimulation. This visual stimulation was provided at the same time points in this control protocol as in the original protocol that included both electrical and visual stimulation (Fig. 1a). Stimulating visually at the same time points provided a control for any changes occurring during the *in vitro* recording due to all factors excluding electrical stimulation. In referring to these cells that were only exposed to visual stimulation, we will use the term visual-only control (VIS-ONLY-CTL, Fig. 1b, right panel).

Consideration of both controls highlights the adaptational effects on visual responses that are separably attributable to electrical stimulation.

### Data Processing and Inclusion Criteria

Commercial spike sorting software was used to process the raw data (Offline Sorter, Plexon Inc., TX, USA). Raw voltage traces (data from both electrical and visual stimulation) were first filtered (using low-cut, 12 point Bessel filter at 51 Hz to exclude line noise); then putative events were detected using a threshold crossing method (4 standard deviations below the mean of the amplitude histogram). These events were sorted into clusters with an automated routine (Standard Expectation Maximization) to assign noise events as well as spiking events multiple cells recorded on each electrode into separate units. Finally, as a quality control step, multiple sorting solutions were manually inspected to identify the best solution and to occasionally modify this solution to minimize Type I and Type II errors in the attribution of events to different sources (Sekhar *et al*. 2016). Units were only judged to contain the spike train from a single RGC and considered for the analysis presented here if they had: 1) a clear lock-out period in the ISI histogram and autocorrelogram, 2) absence of a peak in the cross-correlogram between different cells which would indicate that a single spike train had been incorrectly split into two or more units, 3) good separation in principal component space of a biphasic waveform whose shape is typical of extracellularly recorded action potentials, and 4) stability of the waveform shape and firing rate over the entire experiment. Time stamps of these sorted spikes were collected with NeuroExplorer (Plexon Inc., TX, USA) and exported to MATLAB for further analysis. Accordingly, a total of 2078 WT cells (16 retinal halves) and 1880 *rd10* cells (9 retinal halves) were included in our data set (containing cells for the test and internal control conditions). For the external control condition, a total of 366 WT cells (3 retinal halves) and 573 *rd10* cells (3 retinal halves) were included after data processing. A detailed description of the cell count for electrically responsive cells and visually responsive cells for the test and control conditions are provided in **Table 1**.

**Table 1:**
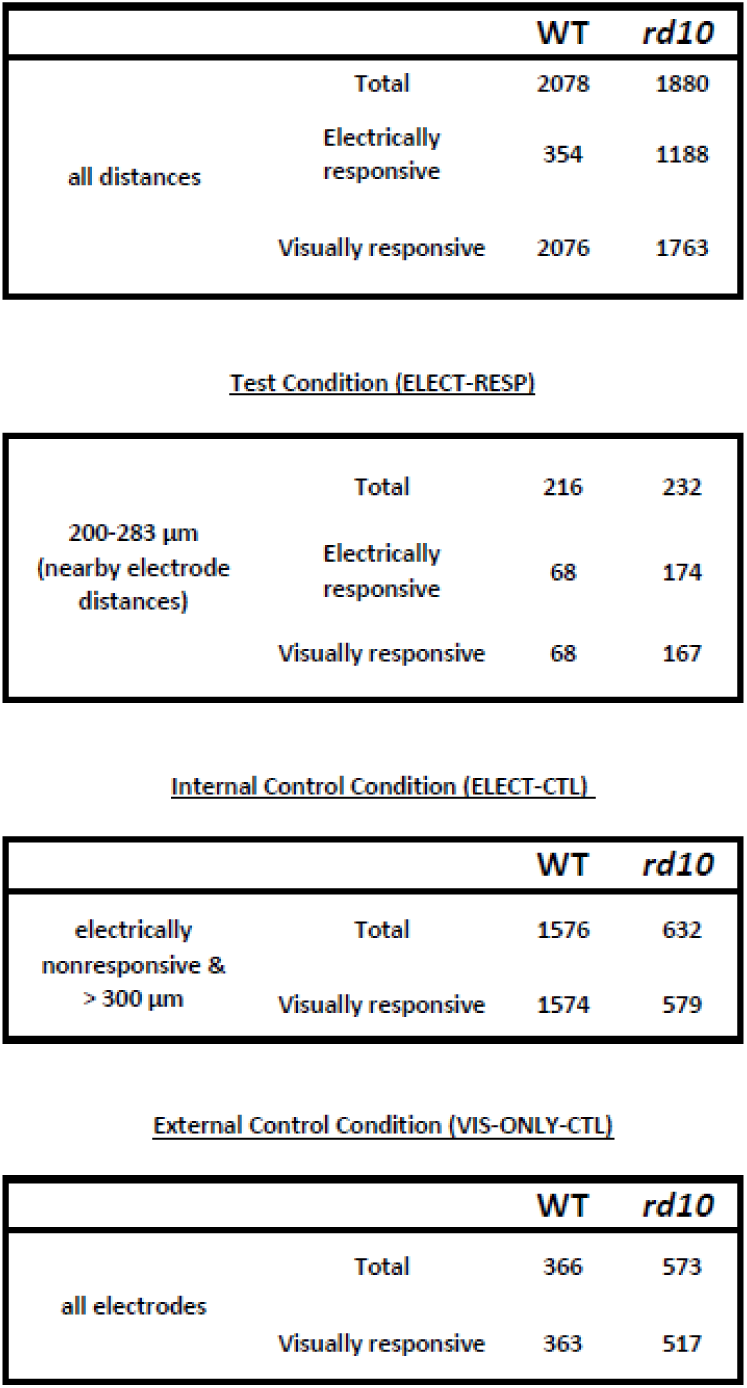
RGC counts for WT and *rd10* for test and control conditions. “distances” are recording distances relative to the stimulating electrode. (For the definition of electrically ‘responsive’ see *Methods: Test and Control Conditions,* and Jalligampala *et al*. 2017*, Methods-Data Analysis*).

|  |  | WT | rd10 |
| --- | --- | --- | --- |
| all distances | Total | 2078 | 1880 |
|  | Electrically responsive | 354 | 1188 |
|  | Visually responsive | 2076 | 1763 |
| Test Condition (ELECT-RESP) |  |  |  |
| 200-283 $\mu\text{m}$<br>(nearby electrode<br>distances) | Total | 216 | 232 |
|  | Electrically responsive | 68 | 174 |
|  | Visually responsive | 68 | 167 |

Internal Control Condition (ELECT-CTL)
|  |  | WT | rd10 |
| --- | --- | --- | --- |
| electrically<br>nonresponsive &<br>> 300 $\mu\text{m}$ | Total | 1576 | 632 |
|  | Visually responsive | 1574 | 579 |

External Control Condition (VIS-ONLY-CTL)
|  |  | WT | rd10 |
| --- | --- | --- | --- |
| all electrodes | Total | 366 | 573 |
|  | Visually responsive | 363 | 517 |

### Data Analysis

#### Determining the visual response parameters

For each visual block of 20 repetitions, the responses were quantified according to the methods of Carcieri *et al*. (2003) (**Fig. 1c**). Briefly, a peristimulus time histogram (PSTH) was generated by aligning each of the 20, 4 s responses, at a 10 ms binning resolution and averaging firing rates across all 20 repetitions. The PSTH was smoothed with a Gaussian filter of σ = 50 ms. To compensate for cells with high firing rates, a baseline firing rate and corresponding SD were calculated for each cell by averaging the firing rates during the last 250 ms of each response phase (ON and OFF, 40 samples per 20-cycle stimulus block). The peak firing rate of each phase (A_1_ and A_2_) was identified to determine time to peak latency (Tp_1_ and Tp_2_) and baseline-subtracted amplitude (A_1_ and A_2_). The duration (a surrogate for response transience) of each response phase (D_1_ and D_2_) was defined as the time over which the baseline-subtracted response falls from peak firing rate (A) to A/*e* (i.e., A_1_ to A_1_/*e* and A_2_ to A_2_/*e*). Amplitude, latency, and duration of a response phase were excluded from analysis if the amplitude did not exceed baseline firing rate + 2SD. Additionally, care was taken to remove any false response peaks arising from sustained responses of one phase extending into the following phase and being detected as a peak (latency < 100 ms). To classify the RGC according to their response polarity, an ON/OFF index was calculated from the amplitudes of ON (A_1_) and OFF (A_2_) responses according to the equation,(A_1_ - A_2_) / (A_1_ + A_2_). Using the ON/OFF index we classified the cells into OFF (−1 to −0.5), ON-OFF (−0.5 to 0.5) and ON (0.5 to 1) categories.

#### Visual response changes

We suspected that electrical stimulation alters visual responses. To test this hypothesis, we compared visual response parameters before and after electrical stimulation (for the test and two control conditions, **Fig. 1d**). At first, we compared the response parameters of the visual block before any electrical stimulation was delivered to the retina to the response parameters of the visual block after the entire electrical stimulation protocol was over. The visual block prior to any electrical stimulation was designated the “**First**” block. The visual block after ∼80 min of electrical and visual stimulation was designated the “**Last**” block. However, for most cells, we observed that the visual response parameters had already changed after only stimulating at lower voltages (0.3 and 0.1 V, ∼20 min) as seen in the example rastergram (**Fig. 1d**). Hence the visual block following 20 min of low voltage electrical stimulation was designated the “**Second**” block and examined for evidence of early visual response parameter changes. It should be noted that although we provided 0.1 V (both cathodic and anodic pulses), this voltage was excluded from the analysis in our previous study due to an uneven current waveform and inconsistent charge delivery (*see Discussion* section *4.1* of Jalligampala *et al*. 2017).

#### Statistics

To test the hypothesis that visual response parameter medians were unchanged between First and Last as well as between First and Second blocks, Wilcoxon’s *ranksum* test was used (MATLAB; The Mathworks, Natick, MA) at a significance of 0.05. To examine whether significant response parameter changes differed between test and each control condition, Wilcoxon’s *ranksum* test, significance of 0.05, was used to test the hypothesis that the two change distributions have equal medians.

## Results

Here, our goal is to gain a deeper understanding of how ongoing electrical stimulation affects the visual response parameters of different RGC types. In a previous study from our group (Jalligampala *et al*. 2017), we established an experimental and analysis framework, by which one could identify the optimal stimulus that will activate a majority of RGCs indirectly via network stimulation from the epiretinal side. This stimulus was optimal for ‘blind’ experiments where the specific response properties for each cell were unknown. During the entire duration of the experiment, six visual stimulus blocks of full-field ‘flash’ stimulus were applied to monitor the stability of RGC responses. These visual blocks were interleaved before, after, and between electrical stimulation blocks spanning ∼80 min of the entire recording time (**Fig. 1a**). Apart from monitoring the stability of the RGC responses, the visual stimulus provided us with an opportunity to classify the cells into different physiological cell types based on their response to the visual stimulus. Surprisingly, we found evidence that visual responses might change over the course of the experiment. In that previous study, we pooled all the RGCs and did not distinguish them into different physiological cell types. However, in this study, we classified RGCs into different response types to better understand how various visual response parameters change during ongoing electrical stimulation. For this study, the primary data set came from the previous study (Jalligampala *et al*. 2017) and was used to test our hypothesis that ongoing electrical stimulation alters visual responses in healthy and degenerating mouse retinas. A preliminary version of this study was reported using only that original data (Jalligampala *et al*. 2015). The additional VIS-ONLY-CTL dataset was collected to account for any non-electrical effects intrinsic to the experimental design.

In an effort to subdivide the RGC population into visual response categories, we examined visual response parameters in the context of a previous study (Carcieri *et al*. 2003). However, we failed to find strong agreement between our data and the statistically determined response distribution boundaries that delineated types in that study (see *Supplement S1, S2*). Nevertheless, due to the predominance of the ON, OFF, and ON-OFF types in the established literature (Morgan & Wong 2007), we divided our data according to the approximate ON/OFF index boundaries of Carcieri *et al*. (see *Methods*).

### Diversity in Alteration of Visual Responses to Electrical Stimulation

To evaluate how electrical stimulation alters the visual response parameters, we plotted the rastergram and peristimulus time histograms (PSTHs) of the cell’s response to full-field flash stimulus (**Fig. 2**). These example cells were both electrically and visually responsive. The rastergram shows the visual response to all six flash blocks presented before, after, and between the electrical stimulation blocks. The PSTH (above) represents the average response (20 trials) of the ‘First block,’ i.e., before electrical stimulation. The PSTH (below) shows the average response (20 trials) of the ‘Last block,’ i.e. after the entire electrical stimulation protocol was over. We observed diversity in the alteration of the visual responses. For our study, our definition of neural adaptation was a change in the cell’s firing rate. Therefore, to evaluate the effect of electrical stimulation, we observed how the cell’s firing rate changed following a period of electrical stimulation. Apart from the changes in firing rate, shown in all the example cells (**Fig. 2a-d**), we also observed changes in other visual response parameters (latency, duration, and ON/OFF index). For some cells, we observed that the latency of the responses became shorter following electrical stimulation (**Fig. 2d**, ON response). For some cells, we observed a change in the transiency of the responses, e.g. some cells which were transient before electrical stimulation became sustained following electrical stimulation (**Fig. 2c, d**). Interestingly, we observed for some cells, a change in ON/OFF index, i.e., before electrical stimulation the cell responded to a single phase of the flash stimulus, however after electrical stimulation the cell responded to both phases of the flash stimulus (**Fig. 2d**, ON to ON-OFF). Such diversified response changes were also observed during the VIS-ONLY-CTL condition in the absence of intervening electrical stimulation.

**Figure 2:**
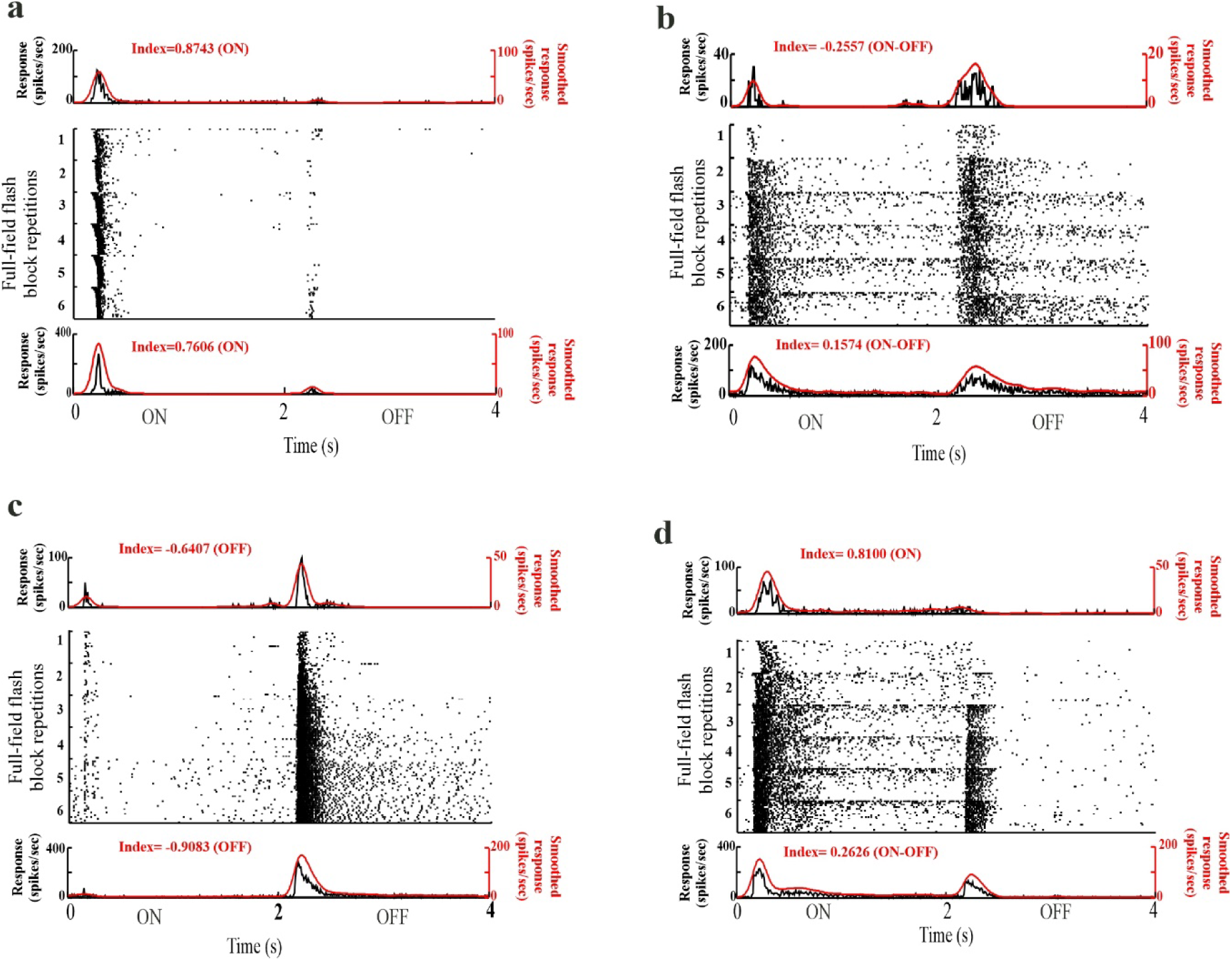
Diversity of visual response changes. **(a-d)** Rastergram depicts responses for all six visual blocks. Response differences between blocks 1 (First), 2 (Second), and 6 (Last) were examined. *(top)* Average PSTH for First block. *(bottom)* Average PSTH for Last block.

As stated, our primary measure of alteration of visual responses (adaptation) in response to electrical stimulation was the change in the cell’s firing rate as quantified in the ON and OFF response amplitudes. Apart from these changes in firing rate, we also measured changes in latency and duration of visual responses. We represent each of these response parameters in notched box-whisker plots (plotting the median, 95% confidence interval of the median, quartiles, and outlier cutoffs) to show the variation across various comparisons (**Fig. 3-5**). We began by examining the changes between the first block and the last block. However, because we observed these changes in visual response parameters as early as ∼20 min (Second block) into the recording (i.e., after the two electrical stimulation blocks of 0.1 and 0.3 V), we also examined the changes of visual response parameters from the first block to the second block. Finally, we examined the changes of the visual responses from the second block to the last block (∼60 min time difference) to understand the slower adaptational effects. We compared these changes between the test condition (ELECT-RESP) and control conditions (ELECT-CTL and VIS-ONLY-CTL; see *Methods* for definitions of test and control conditions). The statistical comparisons are presented in **Table 2**. The cell numbers for each group are available in the *Supplement* (*S6*). To see the outliers of each box plot, refer to the *Supplement* (*S3-5*).

**Figure 3:**
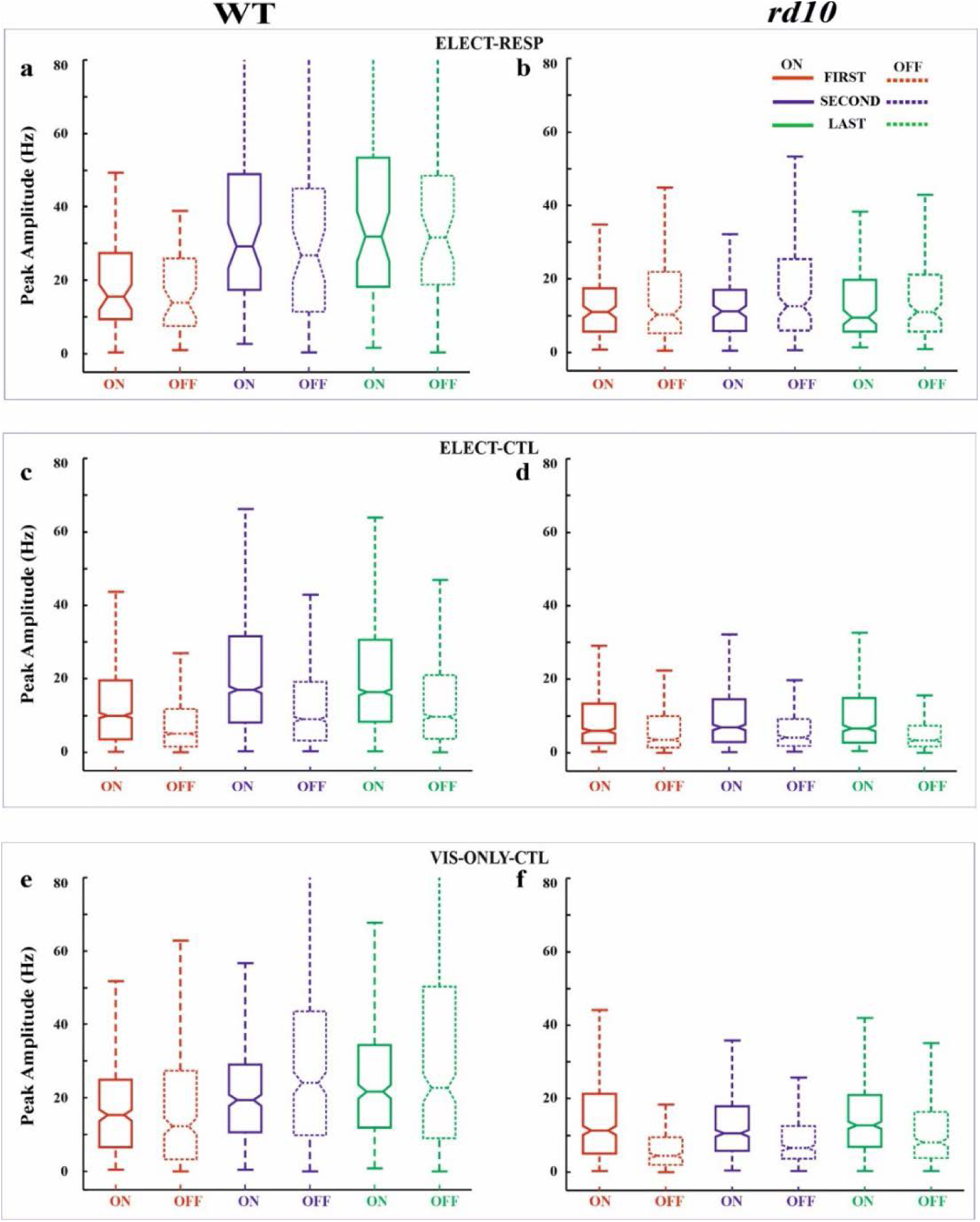
Box-whisker plots for response amplitude. Box plots for ON (solid line) and OFF (dashed) responses shown for First (red), Second (blue), and Last (green) flash blocks for WT **(a, c, e)** and *rd10* **(b, d, f)** retinas for the test (ELECT-RESP, **a**, **b**) and control (ELECT-CTL, **c**, **d** and VIS-ONLY-CTL, **e**, **f**) conditions. Horizontal lines of the box plot demarcate 25%, 50%, and 75% quartiles. Notches are 95% confidence intervals for the median (50% quartile). Whiskers denote data range excluding outliers. Some outlier cutoffs are clipped to show detail. Refer to **Table 2** for pairwise statistical tests.

**Figure 4:**
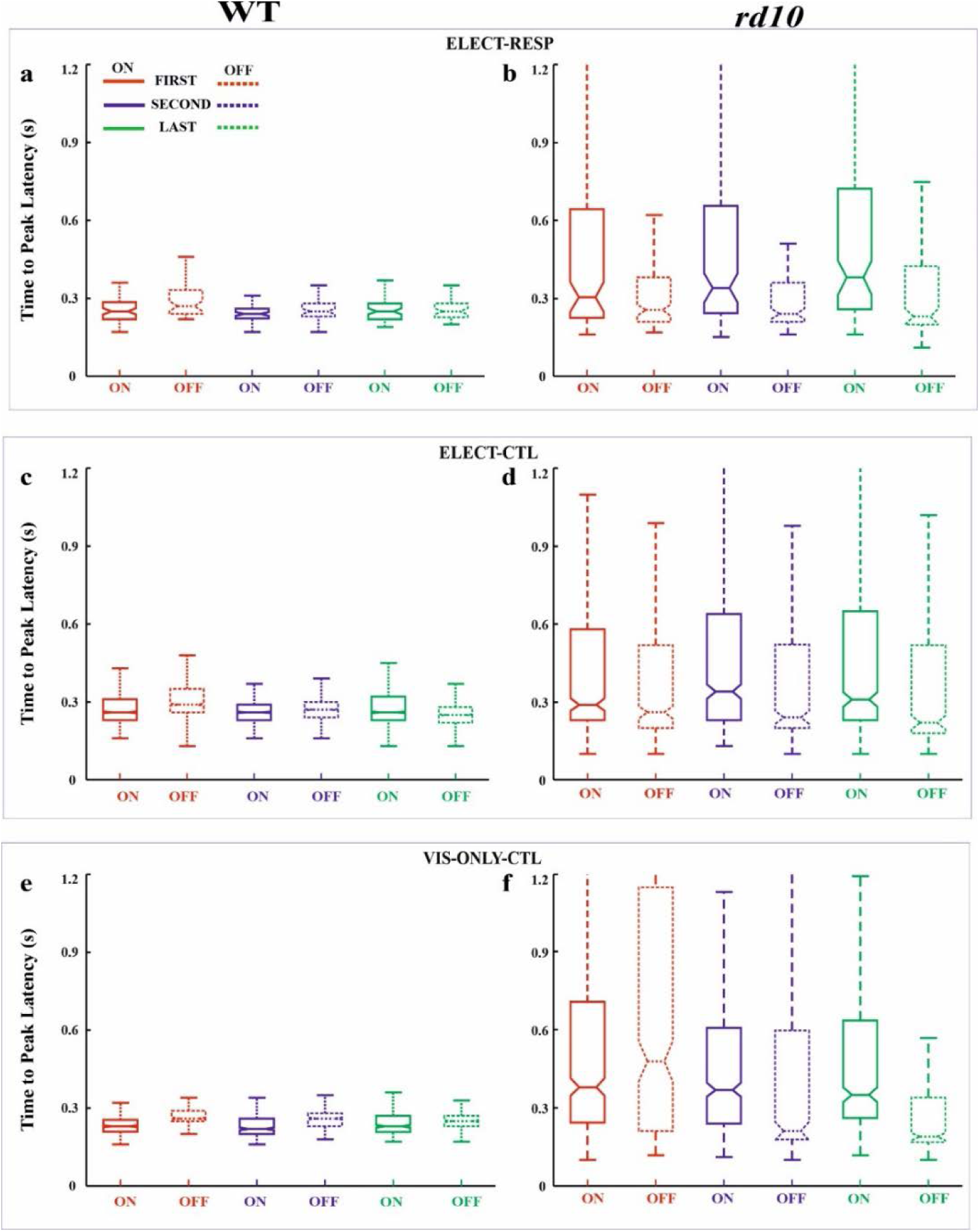
Box-whisker plots for response latency. As in Figure 3.

**Figure 5:**
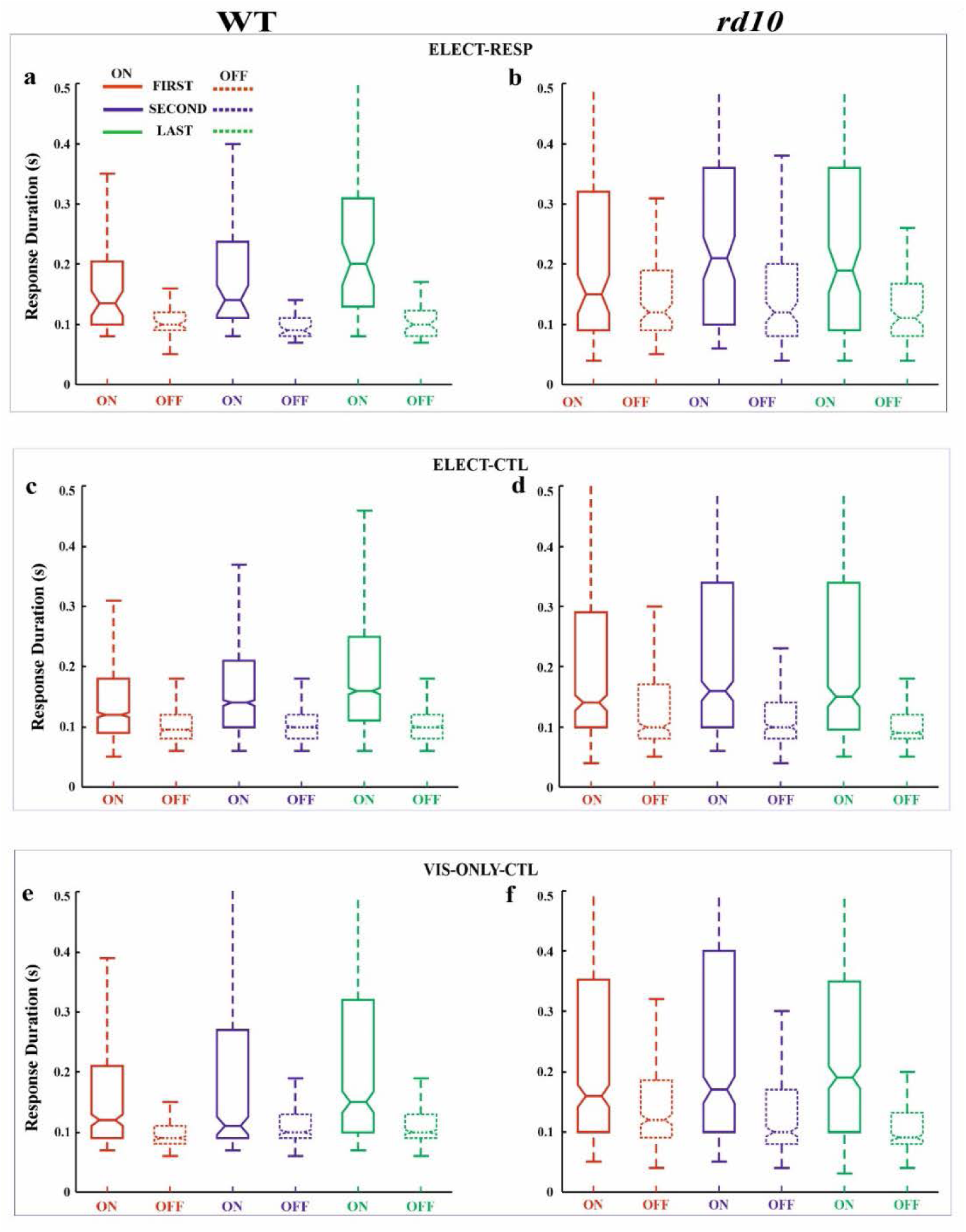
Box-whisker plots for response duration. As in Figure 3.

**Table 2:**
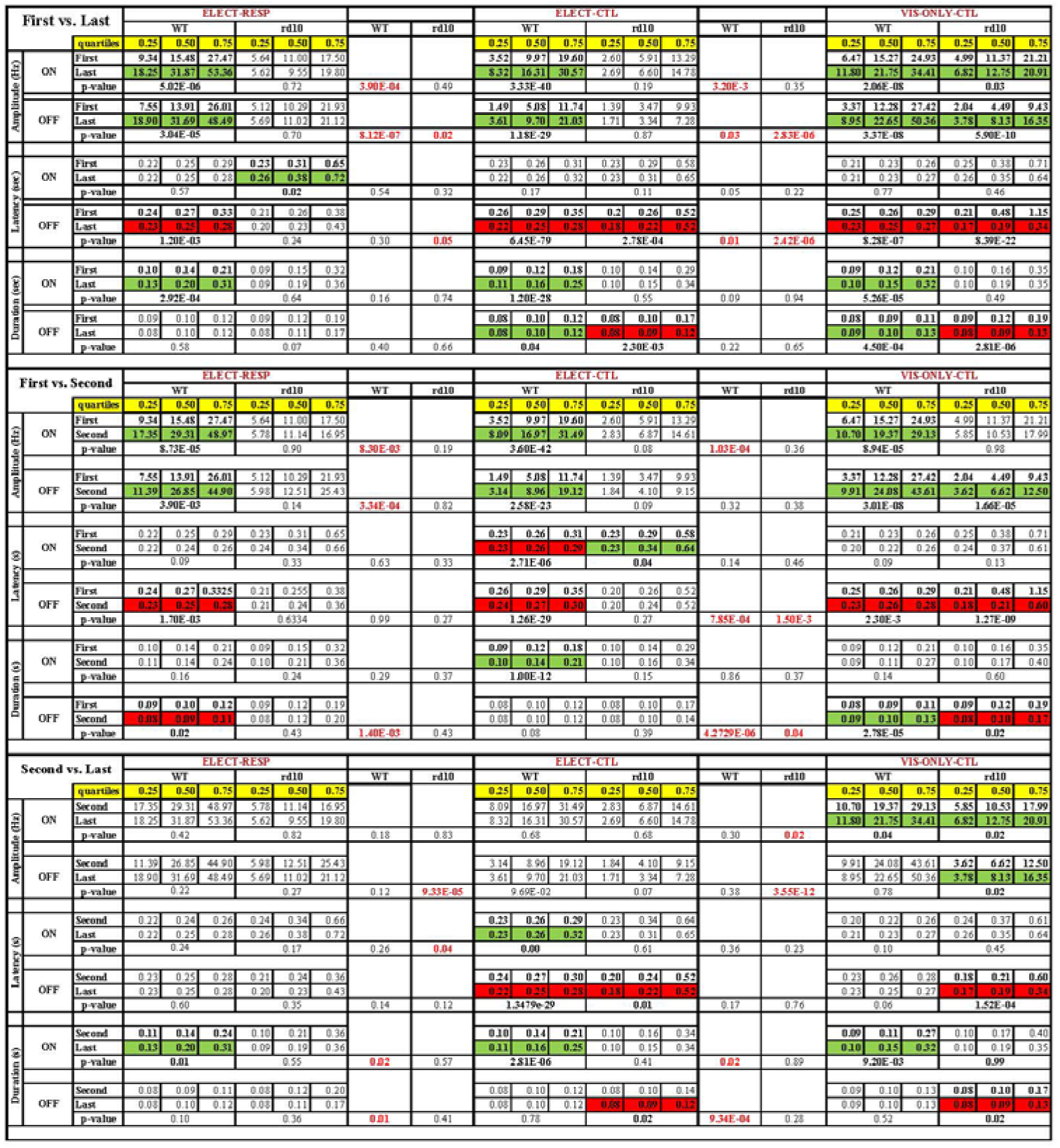
For the plots in **Figures 3-5**, the nonparametric Wilcoxon’s *ranksum* test (MATLAB, p < 0.05) was used to determine significant changes. For each condition (responsive and nearby ELECT-RESP, nonresponsive and distant ELECT-CTL, and the visual-only control VIS-ONLY-CTL) and each mouse strain (WT and *rd10*) we tested whether each visual response parameter differed between First vs. Second, First vs. Last, and Second vs. Last flash stimulus blocks. For each of these tests, the quartiles of the First, Second, and Last response parameter distributions are provided along with the p-value of the test. For testing whether the parameter changes were significantly different between test and control conditions, p-values are presented before the middle and right data blocks. Green boxes identify significant increase. Red boxes identify significant decreases. Bold p-values highlight significant changes. Red p-values indicate significance between test and control conditions.

The main results of our study are as follows. For all three conditions, the firing rate of WT cells increased in response to both light onset and offset, as did the duration of response to light onset. While amplitude increases occurred shortly after onset of electrical stimulation, duration only increased significantly later in the experiment. Critically, these increases were greater for cells that were responsive to electrical stimulation. Secondarily, such adaptation is not consistently observed for *rd10* retina. Many other isolated changes in response latency and duration were observed, but failed to show a large, consistent effect of electrical stimulation. Interestingly, in *rd10* retina under control conditions, the latency and duration of light offset responses significantly decreased over the course of the experiment, although this change was not consistently significant relative to the electrical stimulation test condition. A detailed examination of the observed adaptational changes follows.

#### WT Retinas

##### (1) First vs. Last Block

Amplitude: For both ON and OFF response, there was a significant increase (across all quartiles) in the relative amplitude from the First block to the Last block for all the three conditions (test and both the controls, **Fig. 3a,c,e**). Furthermore, when comparing the magnitude of pre- and post-stimulation changes between the test (ELECT-RESP) and the control conditions (ELECT-CTL and VIS-ONLY-CTL), we found the increase in ON and OFF amplitudes to be significantly greater for the test condition (**Table 2**, green highlights increased values, red p-values are significant). Thus, while response amplitudes increase over time with visual stimulation, this increase is greater if electrical stimulation is also present during that time. Latency: For the latency of ON responses we did not observe any statistical difference between the First and the Last block for all three conditions. The latency of OFF responses significantly decreased for all three conditions (red highlights decreased values). Furthermore, when comparing the magnitude of pre- and post-stimulation changes between ELECT-RESP and VIS-ONLY-CTL, we found the decrease in OFF response latency to be significantly greater for ELECT-RESP, although small in magnitude. There was no statistical difference in latency of OFF response between ELECT-RESP and ELECT-CTL (**Fig. 4a,c,e**). Duration: For all three conditions, there was a significant increase in duration of ON responses, from the First to the Last block. However, when comparing between ELECT-RESP and control conditions, these changes did not differ significantly **(Fig. 5a,c,e)**. Likewise, for the duration of OFF responses, we saw a significant increase from the First to Last block only for the control conditions. When compared to the test condition there was no significant effect of electrical stimulation.

##### (2) First vs. Second Block

Amplitude: For both ON and OFF responses, there was a significant increase in the relative amplitude from the First to the Second block for all three conditions. When comparing the magnitude of pre- and post-stimulation changes between the test and control conditions, we found the increase in ON amplitudes to be significantly greater for ELECT-RESP. However, the increase in OFF response amplitudes was only significantly greater for ELECT-RESP in comparison to the ELECT-CTL, but not (VIS-ONLY-CTL(**Fig. 3a,c,e**). Latency: The latency of the ON responses significantly decreased from the First to the Second block only for ELECT-CTL, and this change was not significantly different when compared to ELECT-RESP (**Fig. 4a,c,e**). For the OFF response latency, there was a significant decrease in latency from the First to the Second block in all conditions. On comparing the test and control conditions, this decrease was significantly greater for ELECT-RESP compared to VIS-ONLY-CTL, but not ELECT-CTL. Duration: There was a significant increase in duration of the ON responses from the First to the Second block only for ELECT-CTL. For the ELECT-RESP, there was a significant decrease in duration of the OFF responses from the First to the Second block, and this change was significant compared to the unchanged OFF duration of ELECT-CTL and the increased OFF duration of VIS-ONLY-CTL (**Fig. 5a,c,e**).

##### (3) Second vs. Last Block

Amplitude; The only amplitude change from the Second to the Last block was for the ON response amplitude of VIS-ONLY-CTL. However, comparing this to ELECT-RESP, there was no statistical significance (**Fig. 3a,c,e**). Latency: Only the ELECT-CTL condition exhibited latency changes from Second to Last block. While the ON response latency increased, the OFF latency decreased (**Fig. 4a,c,e**). Duration: There was a significant increase in duration of the ON responses for all three conditions. Furthermore, comparing the test to the control conditions the magnitude of increase in ON response duration was significantly greater for the ELECT-RESP (**Fig. 5a,c,e**). For all three conditions, there was no significant change in OFF duration from the Second to Last block.

#### rd10 Retinas

##### (1) First vs. Last Block

Amplitude: The only significant amplitude change from First to Last block was an increase in ON and OFF response amplitudes for VIS-ONLY-CTL (**Fig. 3b,d,f**). This increase was only significantly different from ELECT-RESP for the OFF response. Latency: There was a significant increase in ON response latency from the First to Last block for ELECT-RESP. However, this was not a significant change compared to the control conditions. For both control conditions, there was a significant decrease in OFF response latency from the First to the Last block, and this change was also significant relative to the test condition (**Fig. 4b,d,f**). Duration: The only response duration changes from the First to the Last block were a significant decrease in the duration of OFF responses for both control conditions. However, compared to the test condition these changes were not significant (F**ig. 5b,d****,f**).

##### (2) First vs. Second Block

Amplitude: The only change in amplitude from the First to the Second block was for the OFF response of VIS-ONLY-CTL, but this change did not differ from the ELECT-RESP (**Fig. 3b,d,f**). Latency and Duration: For the ON response of ELECT-CTL latency increased from the First to Second block, but was not significant relative to ELECT-RESP. In contrast, both OFF response latency and duration decreased for VIS-ONLY-CTL. While both of these decreases were significant relative to the ELECT-RESP, only the decrease in OFF latency was large in magnitude (**Fig. 4, 5b,d,f**).

##### (3) Second vs. Last Block

Amplitude: The only significant amplitude changes from the Second to the Last block were increases in ON and OFF response amplitudes for VIS-ONLY-CTL. While modest, the magnitude of these changes were significant relative to the ELECT-RESP (**Fig. 3b,d,f**). Latency and Duration: Both latency and duration of the OFF responses for both control conditions decreased from the Second to Last block. Nevertheless, the magnitudes of these changes were small and not significant in comparison to the test condition (**Fig. 4, 5b,d,f**).

### Change in the ON/OFF Index

In **Figure 6** we examined the proportions of RGCs classified as ON, OFF, and ON-OFF before and after electrical stimulation for the test and control conditions. For the WT retinas, we observed a change in the distribution of ON/OFF index values from the First to the Second block for all three conditions. This change in ratio was primarily the change from the purely ON (0.5 to 1) to ON-OFF (−0.5 to 0.5) responses. Given that this change in ON/OFF index was similar for all three conditions, we surmise that this change in weighting of the ON and OFF responses are primarily caused by adaptation of the WT retina to visual stimulus. Furthermore, such adaptation is rather fast as seen by the change from the First to the Second block, but not to the Last block for all the conditions (**Fig. 6a-c**). Interestingly, for the *rd10* retinas, the above observation (ON to ON-OFF) was observed only in the absence of electrical stimulation (VIS-ONLY-CTL, **Fig. 6f**). For both the ELECT-RESP and the ELECT-CTL conditions we saw no change in the ON/OFF index distribution between the three visual blocks (**Fig. 6d, e**). This might suggest that the presence of electrical stimulation prevents a change in the ON/OFF index in the *rd10* retinas regardless of whether or not the RGCs are electrically responsive. Also of interest for the rd10 retina, we found that the relative proportion of OFF cells was greater for the population of RGCs that were responsive to electrical stimulation (ELECT-RESP), but not for the nonresponsive population (ELECT-CTL). We interpret this to reflect that, in the rd10 retina, the relative sensitivity of OFF and ON cells to electrical stimulation is altered to favor OFF cells. The cell numbers contributing to each visual block for the test and control conditions are presented in the *Supplement* (*S7*).

**Figure 6:**
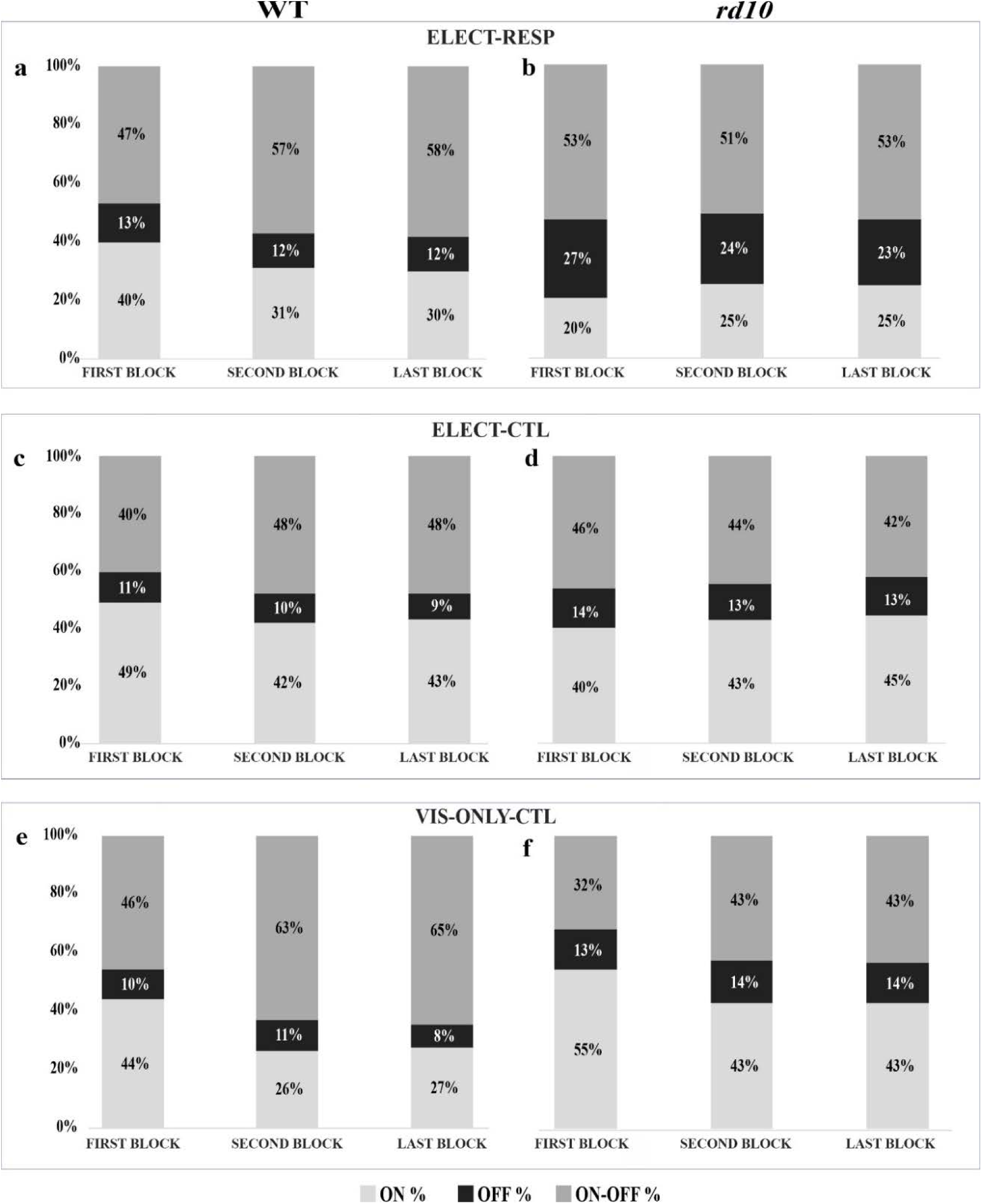
Changes in ON/OFF index. Percentage bar shows the relative proportion of ON, OFF, and ON-OFF responses for First, Second, and Last block, for WT **(a-c)** and *rd10* retina **(d-f)** for all the three conditions (ELECT-RESP, ELECT-CTL, and VIS-ONLY-CTL). Corresponding cell counts are provided in the *Supplement* (*S7*).

## Discussion

In the present study, we focused on discerning the shifts in mouse retinal ganglion cell (RGC) visual response parameters —specifically, latency, duration, and amplitude— before and after electrical stimulation. Our findings illuminate a noteworthy trend: adaptation to visual stimulation *in vitro* involves increased neuronal responsiveness as evidenced by increased firing rate (amplitude) of the cell. Electrical stimulation further strengthens these visual response changes. We are cautious, however, not to draw any further strong conclusions from the changes observed. Given the extensive scope of our analysis, involving nearly 200 statistical comparisons, and the potential for approximately a dozen instances of spurious significance among the 75 observed adaptational and between-condition changes, we recognize the inherent complexity of interpreting such data. For this reason, we have meticulously reported actual p-values and restricted our attention to the most robust adaptation patterns, ensuring a prudent approach to our conclusions.

### Response Changes of Retinal Ganglion Cells in rd10

In retinitis pigmentosa, the loss of photoreceptor cells disrupts the normal functioning of ON and OFF pathways, leading to alterations in adaptation mechanisms. Individuals with RP experience difficulties in adapting to changes in lighting conditions, as well as decreased contrast sensitivity and impaired night vision (Daiger *et al*. 2013, Sahel *et al*. 2015, Sahel *et al*. 2021). This highlights the importance of understanding adaptation mechanisms in retinal degeneration. Research on the adaptation of degenerated mouse retina has revealed several intriguing findings, including functional remodeling, enhanced sensitivity, and plasticity in inner retinal circuits. After the loss of photoreceptor cells, other retinal cell types undergo functional remodeling to compensate for the decreased input from photoreceptors (Jones *et al*. 2016). This remodeling process involves changes in synaptic connectivity, neurotransmitter release, and receptive field properties of retinal neurons. Degenerated retinas often exhibit enhanced sensitivity to dim light stimuli, possibly due to changes in the gain control mechanisms within the retinal circuitry (Stasheff 2008). Lin & Peng have demonstrated plasticity in inner retinal circuits of degenerated retinas, with surviving retinal neurons exhibiting increased dendritic arborization and synaptic remodeling (Lin & Peng 2013). This structural plasticity may contribute to the functional adaptation of degenerated retinas to changes in visual input.

For our rd10 dataset, we observed the majority of adaptational changes for OFF responses. Furthermore, almost all changes were only found in control conditions, suggesting that electrical stimulation might antagonize visual adaptation in the rd10 retina. A previous study has shown that with progressive degeneration in rd10 retinas both the ON and OFF responses are equally affected (Stasheff *et al*. 2011) However, for retinas of the rd1 mouse strain, which is an aggressive form of retinal degeneration, the OFF pathway remains preserved for a longer time span in comparison to the ON pathway (Stasheff 2008). In contrast, even more recent research indicates that the OFF responses are more sensitive to degeneration (Cha *et al*. 2022; Dyszkant *et al*. 2024). Regardless, our observations seem to support the notion that retinal degeneration has differential effects on the ON and OFF pathways and their adapations (Stasheff 2008).

### Role of *in vitro* Adaptation

It is often observed that over the course of long-term *in vitro* recordings the physiological properties of the tissue change (sometimes called ‘expeirmental run-down’). Such changes can lead to changes in visual as well as electrical responses. For example, a recent study showed that increased temperature caused the electrically elicited RGC responses to be more stimulus-locked, with shorter latency and shortened durations (Maturana *et al*. 2015). While such changes could result from changes in pH, temperature, and/or oxygenation, we controlled for these variables in our expeirments. Nevertheless, other variables such as metabolic waste buildup or depletion of cellular resources like glutamate precursors still lead to gradual response changes that eventually conclude with the metabolic death of the tissue. Therefore, to account for such changes during *in vitro* recording, we compared changes associated with electrical stimulation to an external control condition (VIS-ONLY-CTL) in which the same run-down effects and minimal light stimulation (∼4 of 80 minutes) were present.

### Contribution from Visual Adaptation

Work in human ERG (electroretinogram) has shown a gradual increase in amplitude of the a-wave and b-wave components during adaptation to strong light stimulation over a period of approximately 20 min, suggesting that the photoreceptor changes are involved in the rise in amplitudes (Gouras & MacKay 1989). This observation could explain the immediate response changes (∼20 min) observed even in the VIS-ONLY-CTL condition. This increase in response is thought to reflect the redepolarizarion of the cones, after the initial hyperpolarization to an adaptation field. Such redepolarization also restores the horizontal cell polarization. Therefore, apart from any changes induced by the *in vitro* environment, adaptation to visual stimulus also plays a role in the increased response in all three conditions (Webster 2015).

### Characterizing Visual Cell Type Before Electrical Stimulation

We have shown in this study that electrical stimulation can alter visual responses. Therefore, when visual stimulation is to be used for characterizing RGC type, it is strongly recommended that such characterization is performed before any electrical stimulation has been presented. Such characterization is most relevant in healthy retinas or early-stage degenerating retinas, as it is tough to classify the cells during the late stages of degeneration when the visual responses are mostly lost. In addition to visual cell typing, patch-clamp or calcium imaging methods can identify individual cell types morphologically and electrophysiologically. However, it remains untested how electrical stimulation might influence such methods. Recent work from our group employs a newer method to differentiate dozens of visual response types (Shabani *et al*. 2024). Notably, this work also shows signs of response changes following electrical stimulation (see below).

### ON/OFF Index Changes in healthy and rd10 RGCs

In our study, we observed a change in relative weighting for ON and OFF responses, with a shift of purely ON responses to ON-OFF. This shift was mostly observed in WT retinas for all three conditions and in the *rd10* retinas only for the VIS-ONLY-CTL condition. Similar observations have been reported in a previous study in healthy retinas (Tikidji-Hamburyan *et al*. 2015). This prior study showed that, upon full-field stimulation at certain light levels, a cell might be classified as OFF, while at other light levels it would be classified as ON-OFF. Together these observations demonstrate that the relative weighting of ON and OFF response contributions is not fixed and that it can be altered by adaptational mechanisms. In our study, these changes primarily occurred from the First to the Second block (20 min) and remained constant through the Last block (80 min). One possible reason for this observation is that the first visual stimulus is presented after 30 min of no light stimulation (dark adapted state), however, after electrical stimulation and *in vitro* adaptation to periodic light stimulation the retina was in a different adaptational state. For *rd10* retina, however, we did not observe such changes in the test (ELECT-RESP) and internal control (ELECT-CTL). A possible reason may be that the electrical stimulation is antagonistic to visual adaptation in the degenerating retina. Subsequent experiments should examine how visual and electrical adaptation interact in the healthy and degenerating retina at multiple time points during the course of degeneration, although some researchers have already begun to probe this topic (Arens-Arad *et al*. 2020, Arens-Arad *et al*. 2021).

In more recent work from our lab, we have continued to observe visual response changes in the context of a different, ongoing electrical stimulation (Shabani *et al*. 2024). Rather than isolating electrical pulses by 5 seconds, this newer study presented them at a rate of 25 Hz continuously for minutes at a time. The pulse amplitudes spanned a similar range; but were concentrated as a mostly subthreshold Gaussian distribution with mean of −800 mV and SD of 280 mV. Because pulse amplitudes were pulled from this distribution randomly, the stimulus was termed ‘electrical white noise’ – in analogy to similar visual and auditory white noise stimuli that are commonly used in sensory neuroscience research. In Shabani *et al*. 2024, visual response changes were also observed in ON/OFF index, duration, latency, and peak amplitude (**Fig. 7**). The persistence of such visual response adaptation, even for ongoing electrical stimulation that was designed to better match bionic vision implementations, reinforces a need to better understand the relevance of RGC adaptation to bionic vision.

**Figure 7:**
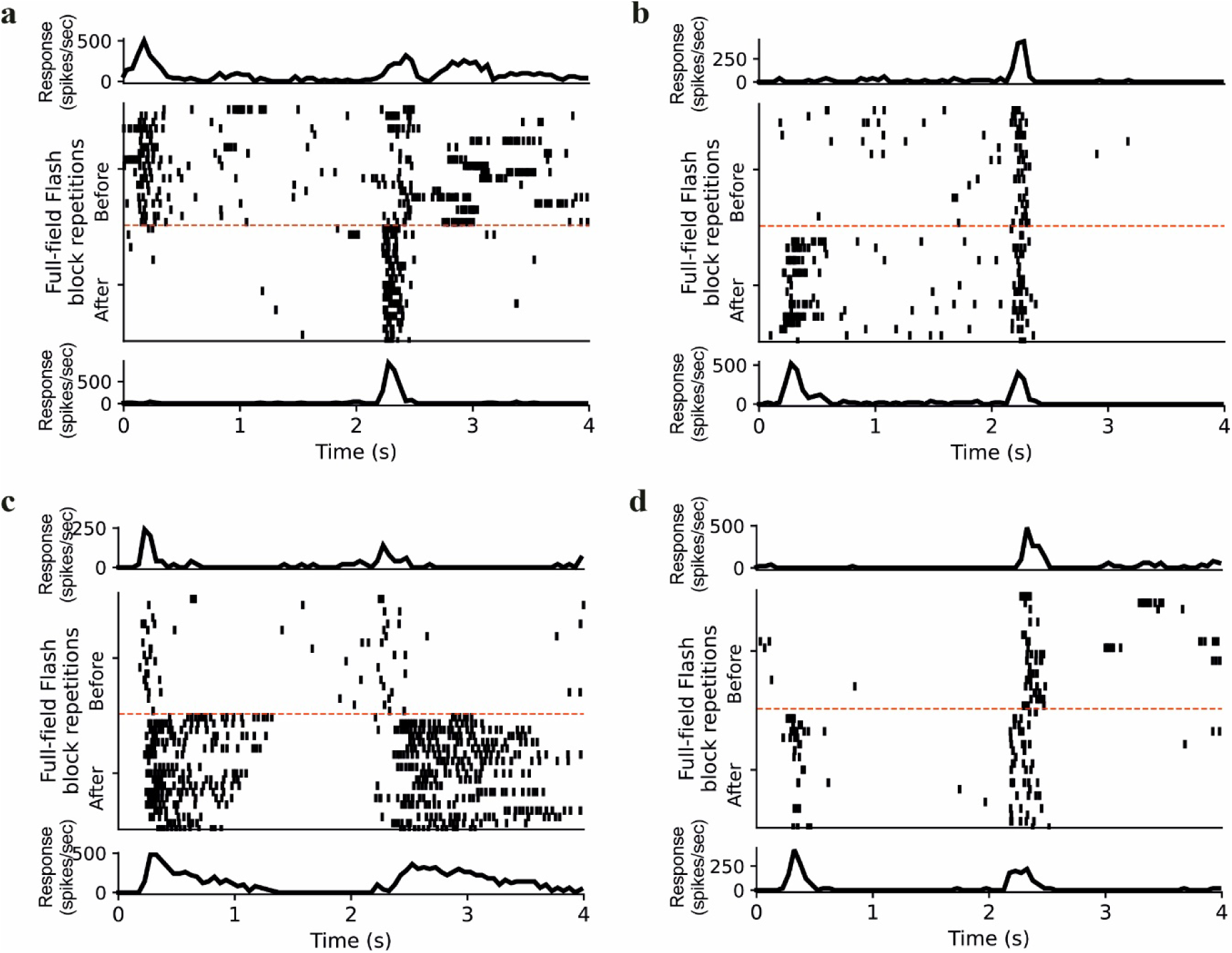
Visual response changes from a separate study. **(a-d)** Rastergram depicts responses for a visual stimulus block before, and one after at least 30 minutes of electrical ‘white noise’ stimulation. *(top)* Average PSTH for block before electrical stimulation. *(bottom)* Average PSTH for block after. Experimental protocol is elaborated in *Methods – Electrical Stimulation* and (Shabani *et al*. 2024).

### Limitations

For the current study, we used a simplified full-field stimulus. With this simplification we could gather responses from thousands of RGCs more quickly, however our ability to unambiguously categorize the visual response of individual cells was diminished. For example, if an ON cell has a strong enough OFF surround, it may be categorized as ON-OFF or even OFF with the present stimulus.

Similarly, response latencies and durations can be contaminated when a full-field stimulus is used. Therefore, using stimulation spots to elicit a response primarily from the visual receptive field center would help in classifying the cell’s response more accurately, but would require streamlined methods for identifying receptive field centers in up to 100 RGCs recorded simultaneously.

This study focused on voltage-controlled stimulation and underlying changes in healthy and rd10 retinas. However, voltage- and current-stimulation may activate different types of retinal neurons in different ways, leading to distinct adaptation profiles. Additionally, the spatial distribution of electrical stimulation may also influence adaptation mechanisms in the retinal network. These issues also warrant further investigation.

### Implications to the Field of Retinal Prostheses

The observed increase in response amplitude is likely the result of alterations in the weighting of network inputs and/or cell-specific physiological properties. Such changes may also affect how the RGCs respond to electrical stimulation of the retinal network. Such effects may have been overlooked in earlier studies since few investigators habituate the retina to ongoing electrical stimulation before examining its electrical responsiveness. An investigation of the cellular and network changes induced by ongoing electrical stimulation – particularly focusing on electrical responsiveness will shed light on this possibility. Given the likelihood that ongoing electrical stimulation will be shown to influence cellular and network responsiveness, and the reality that applied visual prosthetics involve ongoing stimulation, it is advisable to develop paradigms of ongoing background electrical stimulation for future investigations of prosthetic retinal stimulation. One method we have developed is to provide the retina with an ongoing electrical noise stimulus (Sekhar *et al*. 2016). Such stimulation allows electrical responsiveness to be examined in a steady state of adaptation to electrical stimulation that more closely matches the real-world implementation of retinal prosthetic implants.

## Funding & Acknowledgements

This study was supported by the Werner Reichardt Centre for Integrative Neuroscience [CIN] at the Eberhard-Karls University of Tübingen, Excellence Cluster funded by the Deutsche Forschungsgemeinschaft [DFG] within the framework of the Excellence Initiative [**EXC307**, To EZ and **PP 2011-07 & 2013-04** to DLR], the research program of the Bernstein Center for Computational Neuroscience, Tuebingen, funded by the German Federal Ministry of Education and Research [BMBF, **01GQ1002** & **031A308**], the Tistou and Charlotte Kerstan Foundation [to AJ & DLR], the German Ophthalmology Society (DOG) [to AJ], the PRO RETINA Germany foundation for prevention of blindness [to AJ] and European Research Council (ERC) under Grant **101039764** [to ZH].

The authors would especially like to thank Professor Thomas Euler and his laboratory for their guidance and support. The authors gratefully acknowledge the technical assistance of Klaudija Masarini and Norman Rieger. Finally, we thank Professor Shelley Fried for many helpful discussions and his unwavering enthusiasm for this research.

## Competing Financial Interests

None

## Disclosure

Some of the results reported in this article were initially published as a conference proceeding from IEEE NER 2015 (April 20-24), Montpellier, France. This work is reused with permission. Jalligampala A, Zrenner E, Rathbun DL. “Electrical stimulation alters light responses of mouse retinal ganglion cells.” International IEEE/EMBS Conference On Neural Engineering (NER). 2015: 675-678. DOI: 10.1109/NER.2015.7146713

## Author Contributions

AJ & DLR designed the experiments. AJ performed the experiments. AJ, ZH DLR conducted the analysis and interprate data. AJ prepared the manuscript. EZ, ZH& DLR provided critical feedback to the manuscript.

## Supplemental Materials

### Overall Distribution of Response Parameters

Using full-field flash stimulus, we evaluated the distribution of visual response parameters. The response parameters measured were the latency of the response to light on (ON) and light off (OFF), duration of the response to ON and OFF and the relative amplitudes of the response to ON and OFF (*see Data Analysis*). Distributions were examined for both the test (ELECT-RESP) and control conditions (ELECT-CTL and VIS-ONLY-CTL) and both strains of mice (i.e., WT and *rd10*).

#### Test Condition

##### ELECT-RESP: WT

**Latency:** Both ON (**S1 Col 1, Row 1**) and OFF (**S1 Col 2, Row 1**) latencies had a unimodal distribution. Based on the multimodality boundaries for latencies described in Carcieri *et al*. (< 400 ms for short latencies and > 400 ms for long latencies) majority of the cells had short latency response for both flashes of ON and OFF. Duration: The duration of ON response (**S1 Col 3, Row 1**) had a bimodal distribution whereas for the OFF responses (**S1 Col 4, Row 1**) the distribution was rather unimodal. Based on the multimodality boundaries for the duration of responses described in Carcieri *et al*. (< 200 ms for transient cells and > 200 ms for sustained cells) the ON responses had both transient and sustained responses. Whereas, the OFF responses were primarily transient. ON/OFF Index: Based on the relative amplitude of the response to flash ON and flash OFF a bias index was calculated (see *Methods* and Carcieri *et al*. 2003). The distribution of the cells based on the bias index (termed as ON/OFF index for our study) was trimodal dividing the cells into purely ON (+1), purely OFF (−1) and ON-OFF (centered around 0) (**S1 Col 5, Row 1**). The cells which responded to only light onset were classified as purely ON cells. The cells which responded only to light offset were classified as purely OFF cells. Cells which responded to both onset and offset of light were classified as ON-OFF cells. However, based on the distribution, the number of purely ON were higher in comparison to purely OFF cells.

##### ELECT-RESP: rd10

**Latency:** Both ON (**S1 Col 1, Row 2**) and OFF (**S1 Col 1, Row 2**) responses had latencies with multimodal distribution (primarily bimodal) with both short and long latencies. Duration: Duration of both ON (**S1 Col 3, Row 2**) and OFF (**S1 Col 4, Row 2**) responses had a bimodal distribution containing both sustained and transient responses. ON/OFF Index: Similar to the WT retinas, based on the bias index the cells in the *rd10* retina had a trimodal distribution, classifying the cells into purely ON, purely OFF, and ON-OFF cells (**S1 Col 5, Row 2**). In contrast to WT retinas, the number of purely OFF cells were comparatively higher than purely ON cells.

### Control Conditions: (i) Internal Control

#### ELECT-CTL: WT

Latency: Similar to the previous observation of the test condition (ELECT-RESP) both ON and OFF responses had latencies with unimodal distribution, predominantly the short latency response (< 400 ms, **S1 Col 1, 2, Row 3**). Duration: Although the duration of the ON responses had both transient and sustained responses, the distribution was rather unimodal with continuity from transient to sustained responses (< 200 ms transient cells, > 200 ms sustained response, **S1 Col 3, Row 3**). However for the duration of OFF responses were transient with unimodal distribution (**S1 Col 4, Row 3**). ON/OFF Index: Similar to the observation in ELECT-RESP WT cells the distribution based on the bias index was trimodal (**S1 Col 5, Row 3**), classifying the cells in purely ON, purely OFF and purely ON-OFF. Additionally, the number of purely OFF cells were substantially less in comparison to purely ON cells.

#### ELECT-CTL: rd10

Latency: Similar to the previous observation of the test condition (ELECT-RESP) both ON and OFF responses had latencies with multimodal distribution (bimodal) with both short and long latencies (< 400 ms, short and > 400 ms long, **Fig 2. Col 1, 2, Row 4**). Duration: Duration of both ON and OFF responses had a multimodal distribution (bimodal) with both transient and sustained cells (**S1 Col 3, 4, Row 4**). ON/OFF Index: Based on the bias index the distribution of the cells was trimodal (purely ON, purely OFF, ON-OFF). However, unlike the test condition (ELECT-RESP) the number of purely OFF cells were comparatively lower in comparison to purely ON cells (**S1 Col 5, Row 4**).

### Control Conditions: (ii) External Control

#### VIS-ONLY-CTL: WT

Latency: Similar to the test and internal control condition the latency distribution for both ON and OFF responses was unimodal with a short latency (< 400 ms, **S1 Col 1, 2, Row 5**) response. Duration: The distribution of duration of ON responses were bimodal with both transient and sustained responses (**S1 Col 3, Row 5**). For the duration of OFF responses, the distribution was unimodal with transient responses (**S1 Col 4, Row 5**). ON/OFF Index: Similar to the observation for the test and internal control the distribution of the cells was trimodal (purely ON, purely OFF and ON-OFF) and the number of purely OFF cells were comparatively lower to purely ON cells (**S1 Col 5, Row 5**).

#### VIS-ONLY-CTL: rd10

Latency: Similar to previous observations in test and internal control condition the distribution of latencies for ON and OFF response was bimodal with both short and long latency response (**S1 Col 1, 2, Row 6**). Duration: The distribution of duration of both ON and OFF responses was multimodal (bimodal) with both transient and sustained responses (**S1 Col 3, 4, Row 6**). ON/OFF index: Similar to our previous observation in the internal control condition the distribution based on the bias index was trimodal (purely ON, purely OFF, and ON-OFF). The number of purely OFF cells were comparatively lower than purely ON cells.

For the *rd10* retina, although the OFF responses had latencies with a bimodal distribution, the number of cells with short latencies (< 400 ms) were comparatively higher than the long latency responses (> 400 ms). Additionally, for the distribution of the duration of OFF responses, the number of transient cells (< 200 ms) were higher in comparison to sustained cells (> 200 ms).

It should be noted, that for the overall distribution of response parameters described above (**S1**) we could observe a discrepancy between the number of cells for latency, duration, and the ON/OFF index. As mentioned above (see *Data Analysis*) while examining the visual response parameters (latency and duration) for flash ON and flash OFF we excluded any response which had a peak amplitude at latencies < 100 ms (arising from sustained responses extending from an earlier phase). Additionally, responses with nonsignificant amplitude peaks were also excluded while examining these response parameters. However, while determining the ON/OFF index of the cells, relative response amplitudes from both ON and OFF is considered. Hence all cells which had a response amplitude, either for ON or OFF were included in the cell count. Only cells which neither had a response for light onset or light offset were excluded from the analysis.

### Distribution of Response Parameters Based on ON/OFF Index

Next, we evaluated the distribution of visual response parameters (latency and duration) of the cells classified as ON (+0.5 to 1), OFF (−1 to −0.5), and ON-OFF (−0.5 to 0.5) based on the ON/OFF index (Carcieri *et al*. 2003, Sekhar *et al*. 2017). We evaluated the distribution for the test and control conditions and both strains (WT and *rd10* retina).

#### WT ON

*Latency:* For the test condition (ELECT-RESP) and the internal control condition (ELECT-CTL) the distribution of ON response latency was unimodal (**S2 Col 1, Row 1, 3**) with short latency response (< 400 ms). However, for the external control condition (VIS-ONLY-CTL) the distribution of ON latency was bimodal with both short and long latency responses (> 400 ms, **S2 Col 1, Row 5**). *Duration:* For the test condition and the control conditions (both internal and external controls) the distribution of duration of ON responses was multimodal (primarily bimodal, **S2 Col 1, Row 2, 4, and 6**) with both transient (< 200 ms) and sustained responses (> 200 ms).

#### WT OFF

*Latency:* For the test and the control conditions (both internal and external control conditions) the distribution of OFF response latency was unimodal with short latency responses (**S2 Col 2, Row 1, 3, and 5**). For the internal control condition (ELECT-CTL) we did observe few long latency responses. However, most of the cells had a distribution which was primarily unimodal. It should be noted that we did observe a cell count peak at 0. This peak corresponded to the really short latency response amplitudes (< 100 ms). To show the entire cell count distribution for OFF cells, we included these cells in the histogram. *Duration:* For the test condition the distribution for the duration of OFF responses was unimodal with transient responses (**S2 Col 2, Row 2**). However, for both the control conditions the distribution was multimodal (bimodal) with both transient and sustained responses (> 200 ms) (**S2 Col 2, Row 4, 6**).

#### WT ON-OFF

*Latency:* For both the test and control conditions the distribution of ON-OFF response latency was unimodal with short latency responses (**S2 Col 3, Row 1, 3, and 5**). *Duration:* For both test and control conditions the distribution was multimodal (bimodal) with both transient and sustained responses. (**S2 Col 3, Row 2, 4, and 6**).

#### rd10 ON

*Latency:* For both the test and control conditions the distribution of the latency of ON responses was multimodal (bimodal) with both short and long latency responses (**S2 Col 4, Row 1, 3, and 5**). *Duration:* Similar to the latency distribution, the distribution of the duration of ON responses was multimodal (primarily bimodal) with both transient and sustained responses (**S2 Col 4, Row 2, 4, and 6**). For the control conditions, some of the cells had sustained responses lasting up to 2 s.

#### rd10 OFF

*Latency and Duration:* For both the test and control conditions the distribution was rather obscure (for both latency and duration) with a majority of cells peaked around zero (response amplitude of latency < 100 ms). The remaining cells were distributed at various time scales and had no particular distribution. This suggests that the cells contributing to the overall distribution of OFF latency and OFF duration (**S1**) were primarily the cells with ON-OFF responses **(S2 Col 5, Row 1-6**).

#### *rd10* ON-OFF

*Latency and Duration:* For both the test and control conditions the distribution of the ON-OFF response latency was multimodal (primarily bimodal) with both short and long latency responses. For all three conditions we did observe cells which had long latencies > 1 s and in some cells a latency up to 2 s was observed (**S2 Col 6, Row 1, 3, and 5**). Likewise, the distribution of duration of ON-OFF responses was bimodal for both the test and control conditions with both transient and sustained responses, with few cells having a sustained response up to 2 s (**S2 Col 6, Row 2, 4, and 6**)

### Multimodality - Cell Classification

To evaluate if our data set could naturally divide into more than one class, we examined our data set for multimodality. While for the majority of visual responses in WT retina we observed a unimodal distribution, for the *rd10* retina we observed a rather multimodal distribution. This was not surprising, as the age (P28-P37) of the degenerating *rd10* mouse considered in our study had a substantial amount of viable cone photoreceptors which would contribute to the long latency, sustained responses. However, what was intriguing is that for most of the visual responses for WT retinas the responses had a shorter latency and rather a transient response. As the photoreceptors (both rods and cones) are still intact, it was surprising that we did not observe a substantial amount of long latency response. Additionally, for our data set which showed weak multimodality, the physiological response properties were rather a continuum (Carcieri *et al*. 2003, Rodieck 1998). One reason for this could be the use of full-field stimulus rather than a spot stimulus optimized to the receptive field center of each cell (see *Limitations*, below).

---

**S1:**
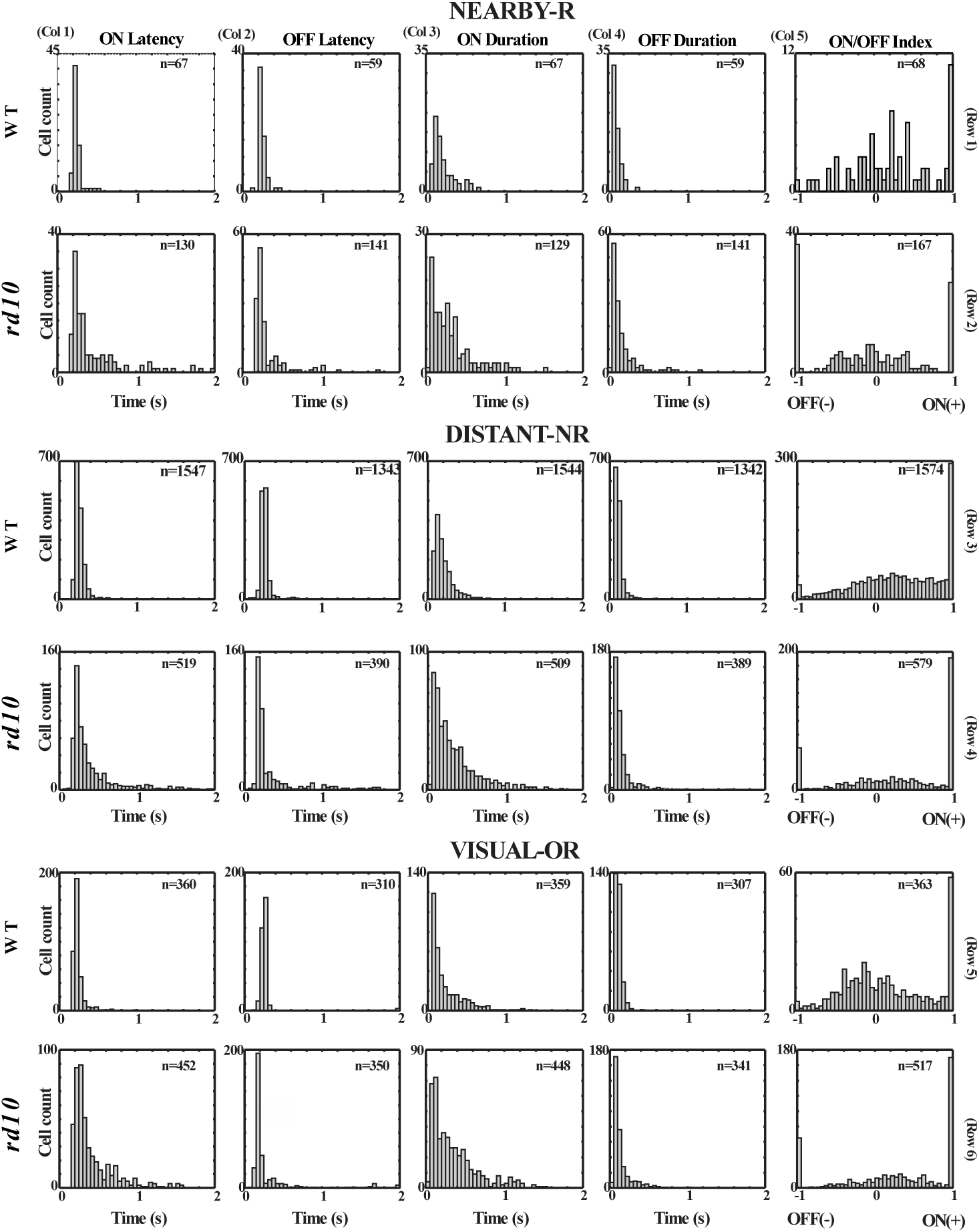
Evaluating multimodality: Overall distribution of response parameters for ON and OFF responses for WT and *rd10* retinas for test (ELECT-RESP) and control conditions (ELECT-CTL and VIS-ONLY-CTL). Responses measured were latency, duration, and relative response amplitudes for computing the ON/OFF index.

**S2:**
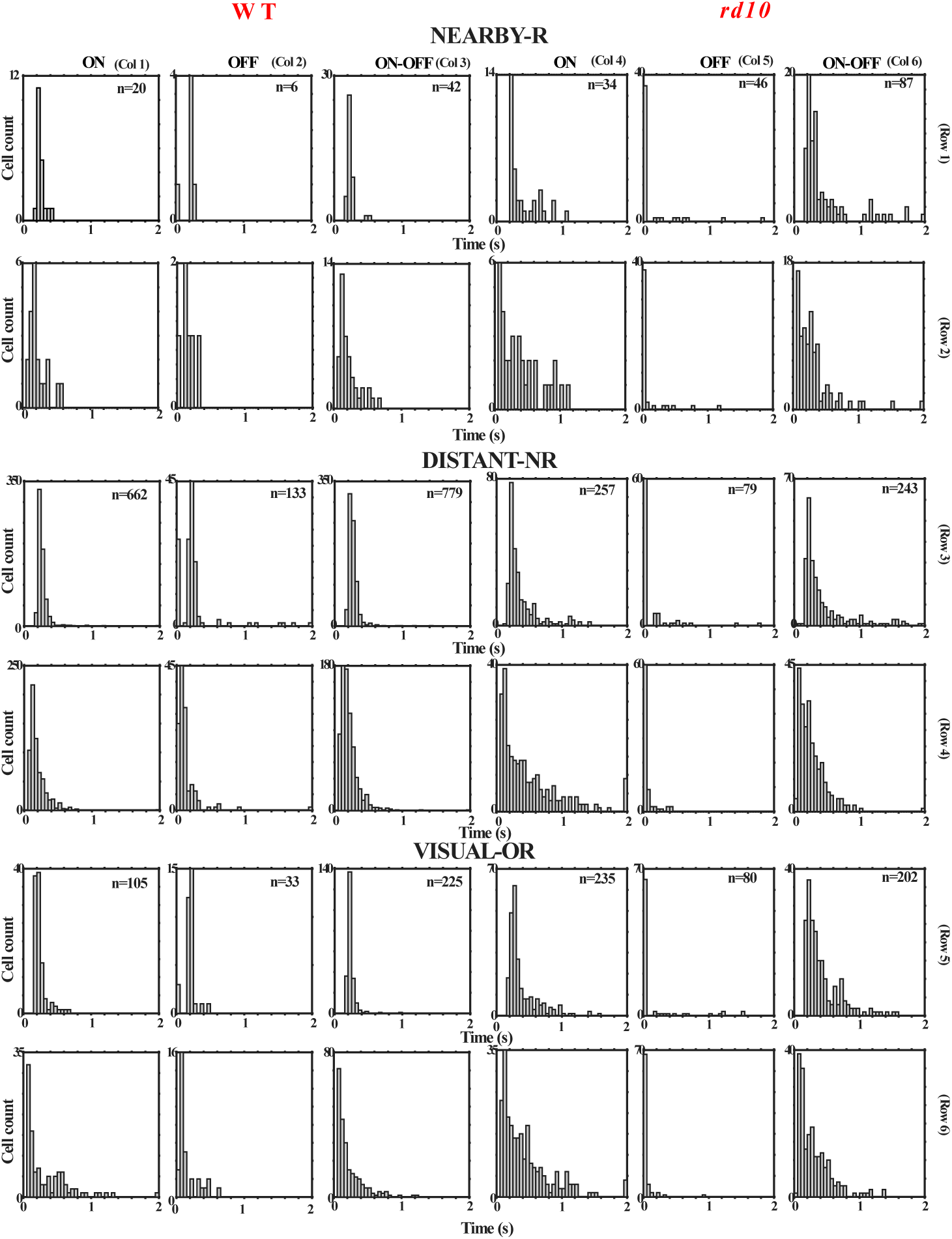
Evaluating multimodality based on ON/OFF index: Overall distribution of latency and duration response parameters of ON, OFF, and ON-OFF RGC types for WT and *rd10* retinas for test (ELECT-RESP) and control conditions (ELECT-CTL and VIS-ONLY-CTL). Only ON response parameters are shown for ON cells, while only OFF parameters are shown for OFF cells. For ON-OFF cells both ON and OFF parameters are shown.

**S3.**
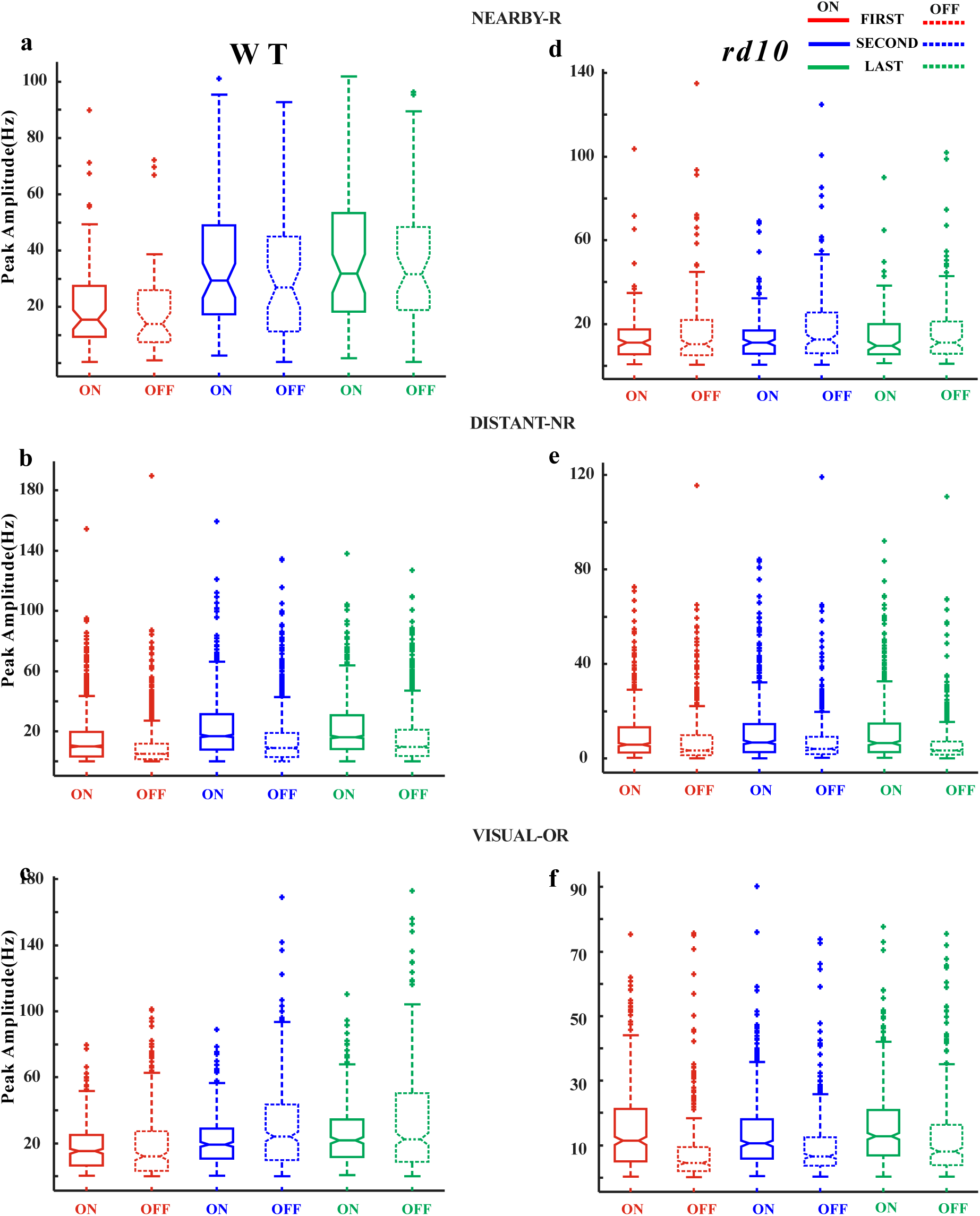
Plotted as in **Figures 3-5**, but including outliers. See **S6** for cell counts of each plot.

**S4.**
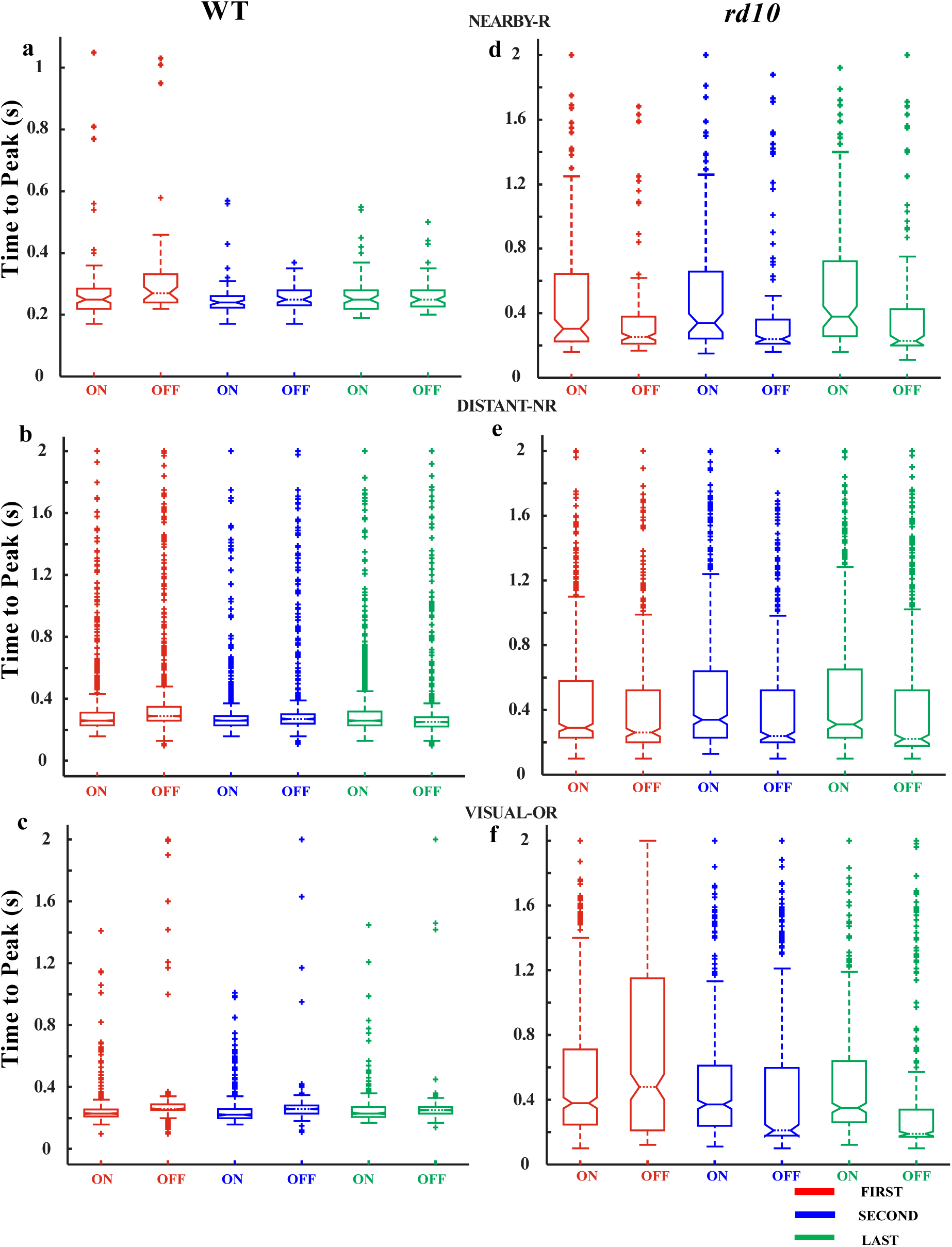
Plotted as in **Figures 3-5**, but including outliers. See **S6** for cell counts of each plot.

**S5.**
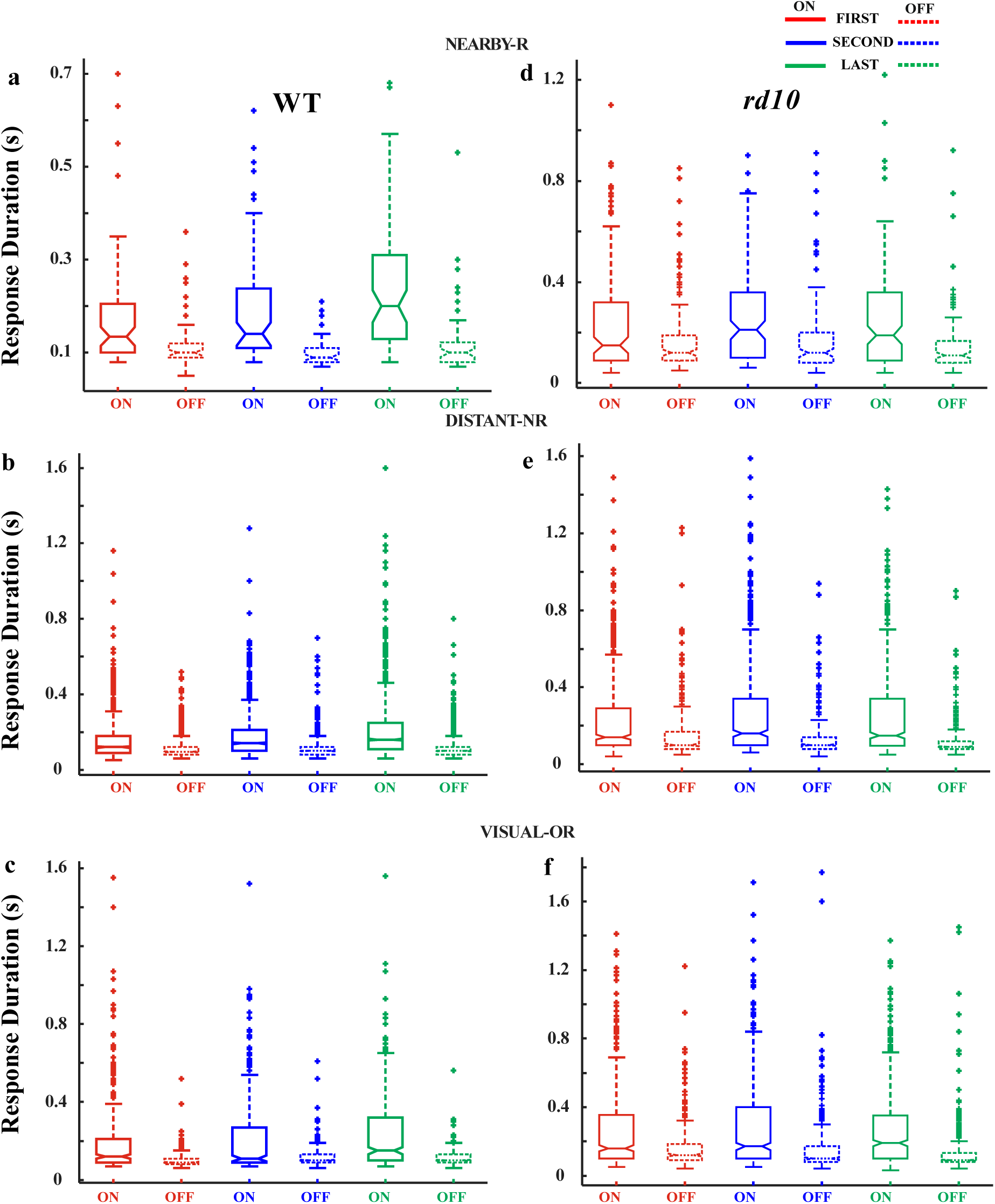
Plotted as in **Figures 3-5**, but including outliers. See **S6** for cell counts of each plot.

**S6:**
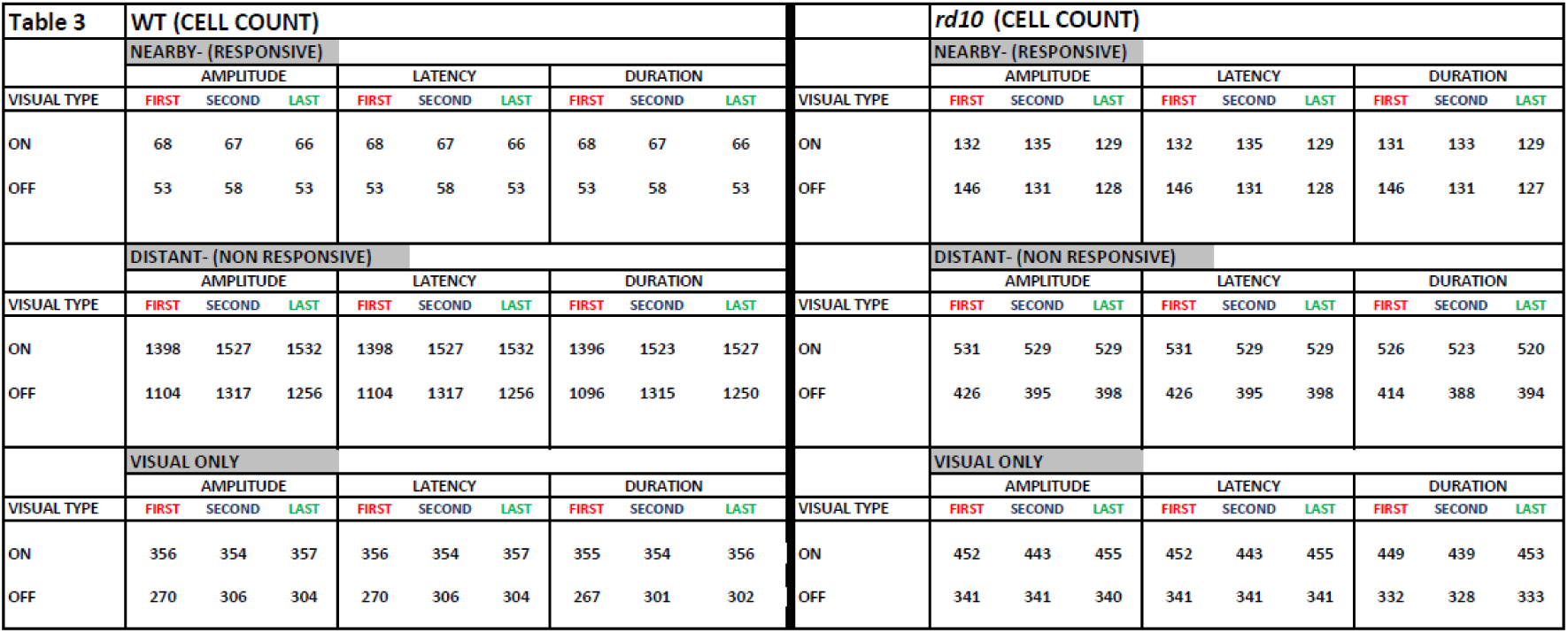
Cell counts for each of the box-whisker plot (for **Fig. 3-5** and **S3-5**).

**S7:**
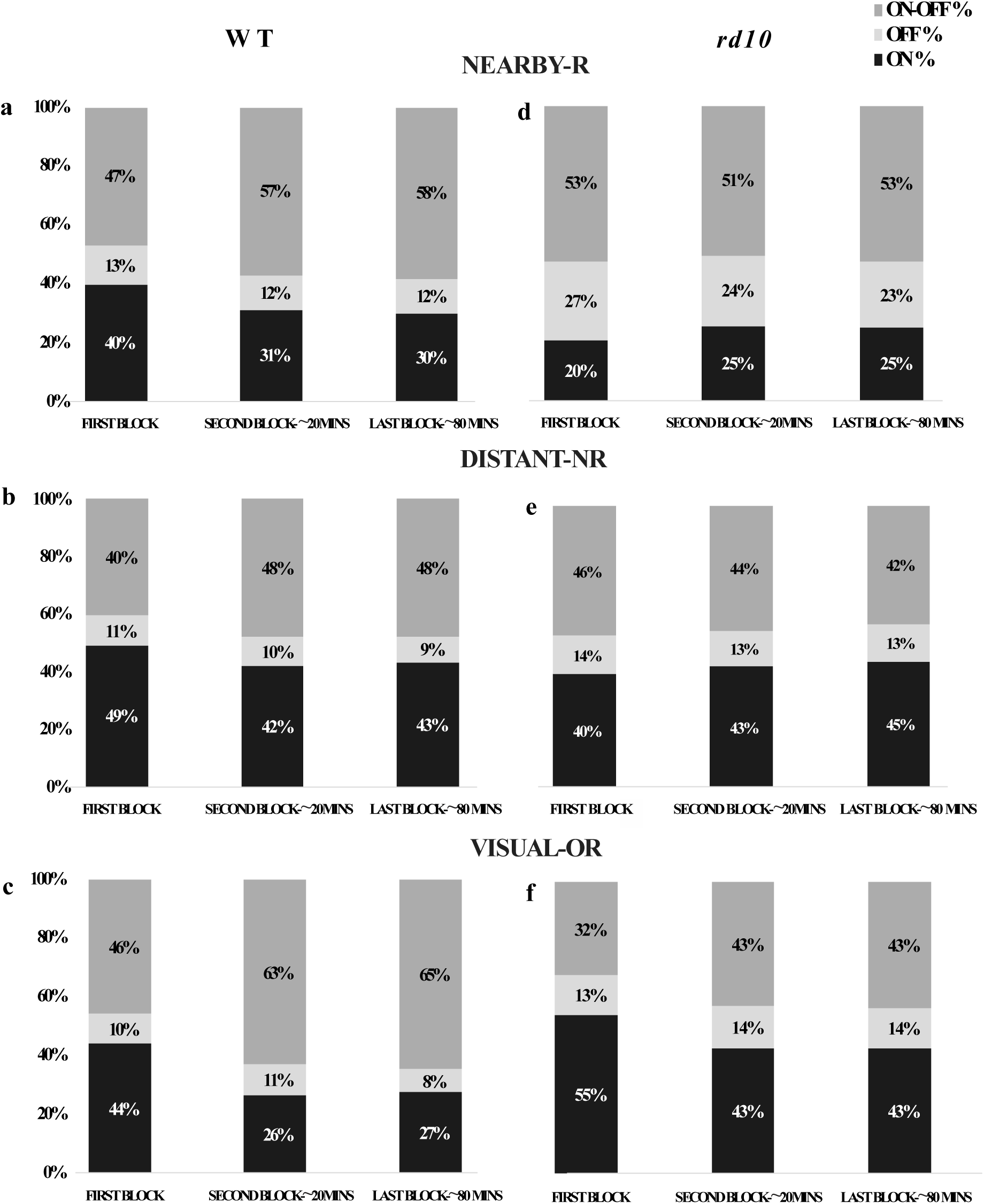
Cell counts for ON, OFF, ON-OFF, and total for evaluating change in ON/OFF index, as shown in **Figure 6**

## Notes

### Competing Interest Statement

The authors have declared no competing interest.

## Bibliography

Arens-Arad T, Farah N, Lender R, Moshkovitz A, Flores T, Palanker D, Mandel Y. Cortical interactions between prosthetic and natural vision. Current Biology. 2020 Jan 6;30(1):176–82.

Arens-Arad T, Lender R, Farah N, Mandel Y. Cortical responses to prosthetic retinal stimulation are significantly affected by the light-adaptive state of the surrounding normal retina. Journal of Neural Engineering. 2021 Mar 3;18(2):026024.

Ayton LN, Barnes N, Dagnelie G, Fujikado T, Goetz G, Hornig R, Jones BW, Muqit MM, Rathbun DL, Stingl K, Weiland JD. An update on retinal prostheses. Clinical Neurophysiology. 2020 Jun 1;131(6):1383–98. DOI: 10.1016/j.clinph.2019.11.029

Baccus SA, Meister M. Fast and slow contrast adaptation in retinal circuitry. Neuron. 2002 Dec 5;36(5):909–19. DOI: 10.1016/s0896-6273(02)01050-4

Baden T, Berens P, Franke K, Román Rosón M, Bethge M, Euler T. The functional diversity of retinal ganglion cells in the mouse. Nature. 2016 Jan 21;529(7586):345–50. DOI: 10.1038/nature16468

Barlow HB. Summation and inhibition in the frog’s retina. The Journal of physiology. 1953 Jan 1;119(1):69. DOI: 10.1113/jphysiol.1953.sp004829

Boinagrov D, Pangratz-Fuehrer S, Goetz G, Palanker D. Selectivity of direct and network-mediated stimulation of the retinal ganglion cells with epi-, sub- and intraretinal electrodes. Journal of neural engineering. 2014 Mar 10;11(2):026008. DOI: 10.1088/1741-2560/11/2/026008

Borghuis BG, Ratliff CP, Smith RG. Impact of light-adaptive mechanisms on mammalian retinal visual encoding at high light levels. Journal of neurophysiology. 2018 Apr 1;119(4):1437–49.

Carcieri SM, Jacobs AL, Nirenberg S. Classification of retinal ganglion cells: a statistical approach. Journal of neurophysiology. 2003 Sep;90(3):1704–13. DOI: 10.1152/jn.00127.2003

Cha S, Ahn J, Jeong Y, Lee YH, Kim HK, Lee D, Yoo Y, Goo YS. Stage-dependent changes of visual function and electrical response of the retina in the rd10 mouse model. Frontiers in Cellular Neuroscience. 2022 Jul 19;16:926096.

Dagnelie G. Retinal implants: emergence of a multidisciplinary field. Current opinion in neurology. 2012 Feb 1;25(1):67–75. DOI: 10.1097/WCO.0b013e32834f02c3.

Daiger SP, Sullivan LS, Bowne SJ. Genes and mutations causing retinitis pigmentosa. Clinical genetics. 2013 Aug;84(2):132–41. DOI: 10.1111/cge.12203

Demb JB. Multiple mechanisms for contrast adaptation in the retina. Neuron. 2002 Dec 5;36(5):781–3. DOI: 10.1016/s0896-6273(02)01100-5

Dyszkant N, Oesterle J, Qiu Y, Harrer M, Schubert T, Gonschorek D, Euler T. Photoreceptor degeneration has heterogeneous effects on functional retinal ganglion cell types. bioRxiv. 2024:2024–09.

Eickenscheidt M, Jenkner M, Thewes R, Fromherz P, Zeck G. Electrical stimulation of retinal neurons in epiretinal and subretinal configuration using a multicapacitor array. Journal of neurophysiology. 2012 May 15;107(10):2742–55. DOI: 10.1152/jn.00909.2011

Fernandez E. Development of visual Neuroprostheses: trends and challenges. Bioelectronic medicine. 2018 Aug 13;4(1):12. DOI: 10.1186/s42234-018-0013-8

Freeman DK, Fried SI. Multiple components of ganglion cell desensitization in response to prosthetic stimulation. Journal of neural engineering. 2011 Jan 19;8(1):016008. DOI: 10.1088/1741-2560/8/1/016008

Fried SI, Hsueh HA, Werblin FS. A method for generating precise temporal patterns of retinal spiking using prosthetic stimulation. Journal of neurophysiology. 2006 Feb;95(2):970–8.

Gouras P, MacKay CJ. Growth in amplitude of the human cone electroretinogram with light adaptation. Investigative ophthalmology & visual science. 1989 Apr 1;30(4):625–30. https://iovs.arvojournals.org/article.aspx?articleid=2160385

Hartline HK. The response of single optic nerve fibers of the vertebrate eye to illumination of the retina. American Journal of Physiology-Legacy Content. 1938 Jan 31;121(2):400–15. DOI: 10.1152/ajplegacy.1938.121.2.400

Ho E, Smith R, Goetz G, Lei X, Galambos L, Kamins TI, Harris J, Mathieson K, Palanker D, Sher A. Spatiotemporal characteristics of retinal response to network-mediated photovoltaic stimulation. Journal of neurophysiology. 2018 Feb 1;119(2):389–400. DOI: 10.1152/jn.00872.2016

Hosseinzadeh Z, Jalligampala A, Zrenner E, Rathbun DL. The spatial extent of epiretinal electrical stimulation in the healthy mouse retina. Neurosignals. 2018 Jan 22;25(1):15–25. DOI: 10.1159/000479459

Hubel DH, Wiesel TN. Receptive fields of single neurones in the cat’s striate cortex. J physiol. 1959 Oct 1;148(3):574–91. DOI: 10.1113/jphysiol.1959.sp006308

Hubel DH, Wiesel TN. Receptive fields, binocular interaction and functional architecture in the cat’s visual cortex. The Journal of physiology. 1962 Jan;160(1):106. DOI: 10.1113/jphysiol.1962.sp006837

Hubel DH, Wiesel TN. Early exploration of the visual cortex. Neuron. 1998 Mar 1;20(3):401–12. DOI: 10.1016/s0896-6273(00)80984-8

Im M, Fried SI. Indirect activation elicits strong correlations between light and electrical responses in ON but not OFF retinal ganglion cells. The Journal of physiology. 2015 Aug 15;593(16):3577–96. DOI: 10.1113/JP270606

Im M, Fried SI. Temporal properties of network-mediated responses to repetitive stimuli are dependent upon retinal ganglion cell type. Journal of neural engineering. 2016 Feb 23;13(2):025002.

Jalligampala A, Sekhar S, Zrenner E, Rathbun DL. Optimal voltage stimulation parameters for network-mediated responses in wild type and rd10 mouse retinal ganglion cells. Journal of neural engineering. 2017 Feb 3;14(2):026004. DOI: 10.1088/1741-2552/14/2/026004

Jalligampala A, Zrenner E, Rathbun DL. “Electrical stimulation alters light responses of mouse retinal ganglion cells.” International IEEE/EMBS Conference On Neural Engineering (NER). 2015: 675–678. DOI: 10.1109/NER.2015.7146713

Jensen RJ, Rizzo JF. Responses of ganglion cells to repetitive electrical stimulation of the retina. Journal of neural engineering. 2007 Jan 24;4(1):S1.

Jones BW, Pfeiffer RL, Ferrell WD, Watt CB, Marmor M, Marc RE. Retinal remodeling in human retinitis pigmentosa. Experimental eye research. 2016 Sep 1;150:149–65. DOI: 10.1016/j.exer.2016.03.018

Kuffler SW. Discharge patterns and functional organization of mammalian retina. Journal of neurophysiology. 1953 Jan 1;16(1):37–68. DOI: 10.1152/jn.1953.16.1.37

Lettvin JY, Maturana HR, McCulloch WS, Pitts WH. What the frog’s eye tells the frog’s brain. Proceedings of the IRE. 1959 Nov;47(11):1940–51.

Lin B, Peng EB. Retinal ganglion cells are resistant to photoreceptor loss in retinal degeneration. PloS one. 2013 Jun 28;8(6):e68084. DOI: 10.1371/journal.pone.0068084

Lorach H, Goetz G, Smith R, Lei X, Mandel Y, Kamins T, Mathieson K, Huie P, Harris J, Sher A, Palanker D. Photovoltaic restoration of sight with high visual acuity. Nature medicine. 2015 May;21(5):476–82.

Maturana MI, Apollo NV, Garrett DJ, Kameneva T, Meffin H, Ibbotson MR, Cloherty SL, Grayden DB. The effects of temperature changes on retinal ganglion cell responses to electrical stimulation. In 2015 37th Annual International Conference of the IEEE Engineering in Medicine and Biology Society (EMBC) 2015 Aug 25 (pp. 7506–7509). IEEE. DOI: 10.1109/EMBC.2015.7320128

Meister M, Pine J, Baylor DA. Multi-neuronal signals from the retina: acquisition and analysis. Journal of neuroscience methods. 1994 Jan 1;51(1):95–106. DOI: 10.1016/0165-0270(94)90030-2

Moore BD, Kiley CW, Sun C, Usrey WM. Rapid plasticity of visual responses in the adult lateral geniculate nucleus. Neuron. 2011 Sep 8;71(5):812–9. DOI: 10.1016/j.neuron.2011.06.025

Morgan J, Wong R. Development of cell types and synaptic connections in the retina. Webvision: the organization of the retina and visual system. 2007 May 29. PMID: 21413410

Multi Channel Systems, 2010. Microelectrode array (MEA) manual, Reutlingen: Multi Channel Systems MCS GmbH.

Olshausen BA, Field DJ. Sparse coding of sensory inputs. Current opinion in neurobiology. 2004 Aug 1;14(4):481–7. DOI: 10.1016/j.conb.2004.07.007

Ray A. Effect of continuous electrical stimulation on retinal structure and function (Doctoral dissertation, University of Southern California). 2010

Rodieck, RW. The first steps in seeing. Sunderland MA (USA): Sinauer Associates; 1998.

Rodieck RW, Stone J. Analysis of receptive fields of cat retinal ganglion cells. Journal of neurophysiology. 1965 Sep 1;28(5):833–49. DOI: 10.1152/jn.1965.28.5.833

Ryu SB, Ye JH, Lee JS, Goo YS, Kim KH. Characterization of retinal ganglion cell activities evoked by temporally patterned electrical stimulation for the development of stimulus encoding strategies for retinal implants. Brain research. 2009 Jun 12;1275:33–42. DOI: 10.1016/j.brainres.2009.03.064

Sahel JA, Boulanger-Scemama E, Pagot C, Arleo A, Galluppi F, Martel JN, Esposti SD, Delaux A, de Saint Aubert JB, de Montleau C, Gutman E. Partial recovery of visual function in a blind patient after optogenetic therapy. Nature medicine. 2021 Jul;27(7):1223–9. DOI: 10.1038/s41591-021-01351-4

Sahel JA, Marazova K, Audo I. Clinical characteristics and current therapies for inherited retinal degenerations. Cold Spring Harbor perspectives in medicine. 2015 Feb 1;5(2):a017111. DOI: 10.1101/cshperspect.a017111

Sekhar S, Jalligampala A, Zrenner E, Rathbun DL. Tickling the retina: integration of subthreshold electrical pulses can activate retinal neurons. Journal of neural engineering. 2016 May 17;13(4):046004. DOI: 10.1088/1741-2560/13/4/046004

Sekhar S, Jalligampala A, Zrenner E, Rathbun DL. Correspondence between visual and electrical input filters of ON and OFF mouse retinal ganglion cells. Journal of neural engineering. 2017 Jun 12;14(4):046017. DOI: 10.1088/1741-2552/aa722c

Shabani H, Sadeghi M, Zrenner E, Rathbun DL, Hosseinzadeh Z. Classification of pseudocalcium visual responses from mouse retinal ganglion cells. Visual Neuroscience. 2021 Jan;38:E016. DOI: 10.1017/S0952523821000158

Shabani H, Zrenner E, Rathbun DL, Hosseinzadeh Z. Electrical Input Filters of Ganglion Cells in Wild Type and Degenerating rd10 Mouse Retina as a Template for Selective Electrical Stimulation. IEEE Transactions on Neural Systems and Rehabilitation Engineering. 2024 Jan 31. DOI: 10.1109/TNSRE.2024.3360890

Sherrington CS. Observations on the scratch-reflex in the spinal dog. The Journal of physiology. 1906 Mar 3;34(1-2):1. DOI: 10.1113/jphysiol.1906.sp001139

Stasheff SF. Emergence of sustained spontaneous hyperactivity and temporary preservation of OFF responses in ganglion cells of the retinal degeneration (rd1) mouse. Journal of neurophysiology. 2008 Mar;99(3):1408–21. DOI: 10.1152/jn.00144.2007

Stasheff SF, Shankar M, Andrews MP. Developmental time course distinguishes changes in spontaneous and light-evoked retinal ganglion cell activity in rd1 and rd10 mice. Journal of neurophysiology. 2011 Jun;105(6):3002–9. DOI: 10.1152/jn.00704.2010

Stett A, Mai A, Herrmann T. Retinal charge sensitivity and spatial discrimination obtainable by subretinal implants: key lessons learned from isolated chicken retina. Journal of neural engineering. 2007 Feb 20;4(1):S7. DOI: 10.1088/1741-2560/4/1/S02

Tikidji-Hamburyan A, Reinhard K, Seitter H, Hovhannisyan A, Procyk CA, Allen AE, Schenk M, Lucas RJ, Münch TA. Retinal output changes qualitatively with every change in ambient illuminance. Nature neuroscience. 2015 Jan;18(1):66–74. DOI: 10.1038/nn.3891

Tikidji-Hamburyan A, Reinhard K, Storchi R, Dietter J, Seitter H, Davis KE, Idrees S, Mutter M, Walmsley L, Bedford RA, Ueffing M. Rods progressively escape saturation to drive visual responses in daylight conditions. Nature communications. 2017 Nov 27;8(1):1813. DOI: 10.1038/s41467-017-01816-6

Weber, E. H. (1846). Tastsinn und Gemeingefuehl. In Rudolph Wagners Handwörterbuch der Physiologie. [Reprinted (1905) in E. Hering (Ed.) Ostwald’s Klassiker der exakten Wissenschaften, No. 149. Leipzig: W. Engelmann]

Webster MA. Visual adaptation. Annual review of vision science. 2015 Nov 24;1(1):547–67. DOI: 10.1146/annurev-vision-082114-035509

Wilke R, Gabel VP, Sachs H, Schmidt KU, Gekeler F, Besch D, Szurman P, Stett A, Wilhelm B, Peters T, Harscher A. Spatial resolution and perception of patterns mediated by a subretinal 16-electrode array in patients blinded by hereditary retinal dystrophies. Investigative ophthalmology & visual science. 2011 Jul 1;52(8):5995–6003. DOI: 10.1167/iovs.10-6946

